# Rpd3L Coordinates Chromatin State and Transcription-Replication Conflict Resolution through H3K4 Methylation-Dependent and Independent Mechanisms

**DOI:** 10.64898/2026.04.13.718173

**Authors:** Shin Yen Chong, Ya-Ling Chen, Yueh-Tzu Hsu, Chia-Ling Hsu, Tsai-Ming Lu, Yi-Chen Lo, Cheng-Fu Kao

## Abstract

Faithful genome duplication requires coordination between transcription and replication. Disruption of this coordination causes transcription–replication conflicts (TRCs), leading to replication stress and genome instability. How chromatin regulators modulate these processes remains unclear. Here, we show that the Rpd3L histone deacetylase complex dynamically modulates chromatin state to control replication fork progression and buffer TRCs in *Saccharomyces cerevisiae*. Rpd3L is targeted through both histone H3 lysine 4 methylation-dependent recruitment and methylation-independent mechanisms engaged under replication stress. Loss of H3K4 methylation or Rpd3L function promotes histone acetylation, accelerates fork progression through transcribed regions, and increases transcription-associated genome instability. Balanced acetylation at multiple histone lysines is required to stabilize replication forks under stress. While histone deacetylase complexes have been implicated in repairing damaged forks, our findings reveal that Rpd3L acts preemptively to modulate chromatin state and replication dynamics during TRCs, defining a chromatin-based mechanism that safeguards genome stability.

## Introduction

Faithful genome duplication requires precise coordination between DNA replication and transcription, two essential processes that share the same DNA template. Disruption of this coordination can give rise to transcription-replication conflicts (TRCs), which stall replication forks, generate DNA breaks, and contribute to genome instability (*1, 2*). In both yeast and metazoans, TRCs are now recognized as a major source of endogenous DNA damage (*3*), and their resolution requires the interplay of replication checkpoint proteins, RNA processing factors, and chromatin remodelers (*4*).

Chromatin modifications such as histone acetylation and methylation are key regulators of DNA accessibility and polymerase engagement (*5, 6*). Among these, methylation of histone H3 on lysine 4 (H3K4me) is broadly associated with active transcription (*7, 8*), but recent work suggests it also plays a role in modulating replication fork dynamics under stress (*9, 10*). In budding yeast, loss of H3K4 methylation suppresses replication fork instability and reduces genome instability in checkpoint-deficient cells, revealing a counterintuitive role for this “transcription-active” mark in impeding fork progression when checkpoint signaling is compromised (*9*). Consistent with this, Set1-dependent H3K4 methylation has been shown to limit DNA damage under replication stress by shaping chromatin environments that protect replication forks from transcription-associated obstacles (*10, 11*). These findings suggest that the epigenetic landscape may serve not only to facilitate transcription but also to shape replication timing and fork stability, especially at transcribed regions.

Histone deacetylases (HDACs) play a central role in regulating chromatin structure by removing acetyl groups from lysine residues, thereby reducing histone mobility and transcriptional accessibility (*12*). In *Saccharomyces cerevisiae*, the Rpd3 HDAC exists in two major complexes: Rpd3L (Large) and Rpd3S (Small) (*13, 14*). Although both share the Rpd3 catalytic subunit, they are structurally and functionally distinct (*15*). Rpd3L, but not Rpd3S, has been implicated in stress responses, gene regulation, and genome maintenance (*16–19*). Rpd3L contains the PHD finger protein Pho23, which can directly bind H3K4me3 (*19, 20*), positioning it as a candidate effector for methylation-sensitive chromatin regulation. Notably, this functional module is evolutionarily conserved: in mammals, the PHD finger protein ING2 similarly binds H3K4me3 and recruits Sin3A-HDAC complexes, linking methylation to transcriptional repression and chromatin remodeling (*21, 22*). Despite its known roles in transcriptional repression, the function of Rpd3L in controlling replication fork behavior remains poorly defined (*18*). In particular, it is unclear whether its recruitment and activity are strictly dependent on H3K4 methylation, or whether it can also function through alternative pathways to regulate chromatin under replication stress.

Here, we reveal an unanticipated function of the Rpd3L histone deacetylase complex in safeguarding genome integrity during transcription-replication conflicts (TRCs) in *Saccharomyces cerevisiae*. Through a focused genetic screen, we identified H3K4 methylation as a key modulator of genome instability under replication stress and uncovered Rpd3L as a downstream effector in this pathway. Integrating genetic analysis with chromatin profiling and replication dynamics assays, we demonstrate that Rpd3L curtails replication fork progression at transcribed loci to limit TRC-associated damage. Notably, this regulation occurs through both H3K4 methylation-dependent and -independent mechanisms, establishing Rpd3L as a central chromatin regulator of the replication stress response.

## Results

### H3K4 methylation and Rpd3L complex coordinate regulation of replication stress response

Our previous genetic studies demonstrated that loss of H3K4 methylation suppresses the replication stress sensitivity of *rad53* mutants in *Saccharomyces cerevisiae*, suggesting that H3K4me contributes to replication fork reversal when checkpoint signaling is compromised (Fig. 1A, top) (*9*). Since H3K4 methylation does not directly alter chromatin accessibility (*5*), its biological effects are likely mediated through specific reader proteins that recognize this mark and recruit downstream effectors (Fig. 1A, left). We hypothesized that reader proteins that interact with H3K4me3 act as functional effectors that modulate replication fork dynamics and mitigate transcription-replication conflicts (TRCs) under stress conditions.

**Fig. 1.**
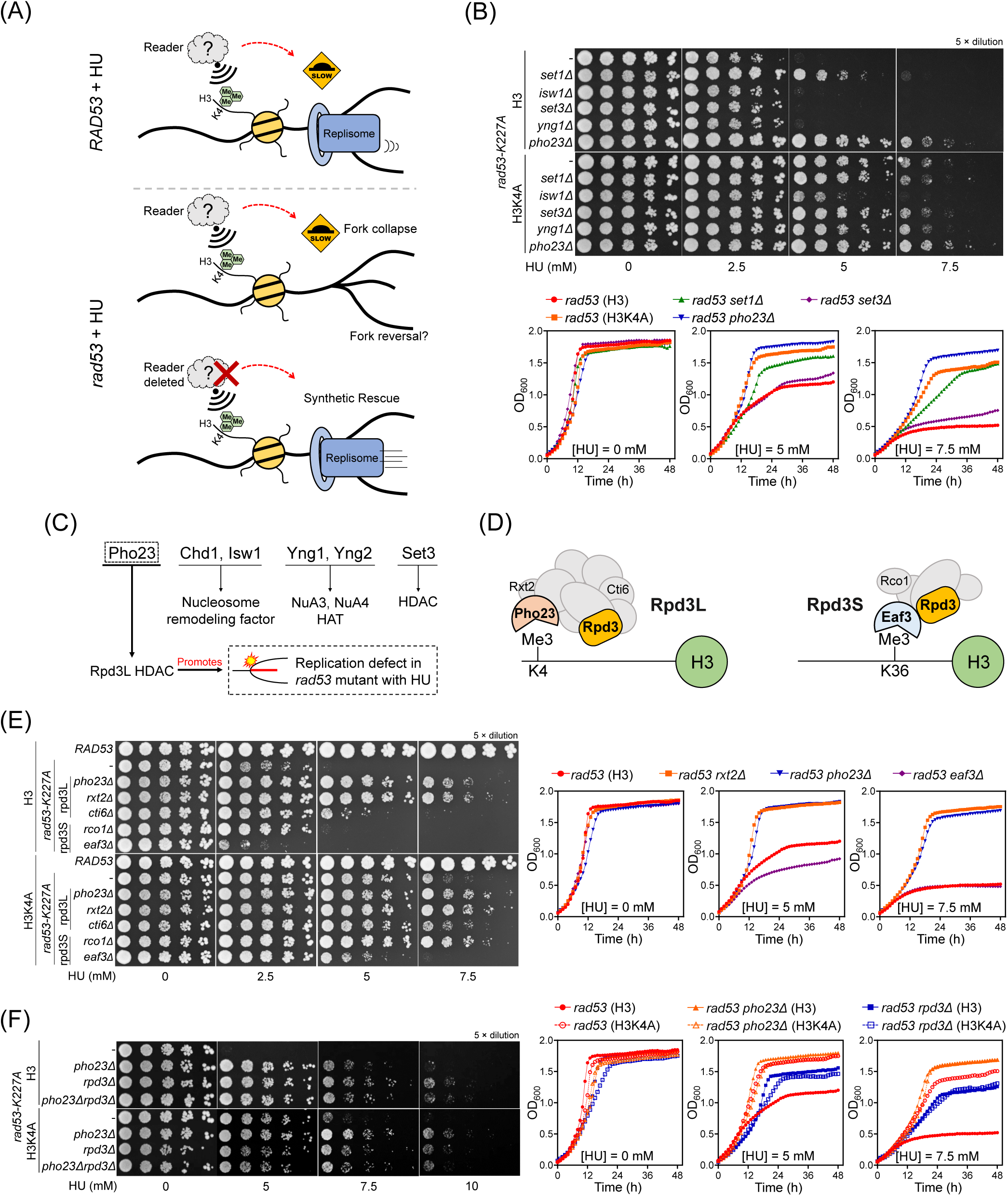
Pho23 and the Rpd3L complex mediate the effects of H3K4 methylation on replication stress sensitivity in *rad53-K227A* mutants. (A) Schematic illustration summarizing the relationship between H3K4 methylation, reader proteins and replication stress sensitivity examined in this study. In *rad53-K227A* cells under hydroxyurea (HU)-induced stress, H3K4me3 impedes fork progression, potentially through recruitment of chromatin effectors such as Pho23. Deletion of specific readers may alleviate replication stress via synthetic rescue. **(B)** Spot dilution assays (upper panel) and growth curves (lower panel) showing HU sensitivity of *rad53-K227A* cells carrying deletions of *SET1*, known H3K4me readers (*ISW1*, *SET3*, *YNG1*, and *PHO23*), or the H3K4A histone mutant. Deletion of *PHO23*, similar to H3K4A or *set1Δ*, suppresses HU sensitivity, whereas deletions of other reader genes show no detectable effect. **(C)** Schematic representation of chromatin complexes containing H3K4me readers. Pho23 associates with the Rpd3L complex, whereas other readers are linked to chromatin remodelers or HAT complexes. **(D)** Schematic representation of selected Rpd3L and Rpd3S subunits with distinct chromatin-targeting properties. **(E)** Spot assays (left panel) and growth curves (right panel) comparing HU sensitivity of *rad53-K227A* strains with deletions of Rpd3L or Rpd3S components, in the presence or absence of H3K4A. Rpd3L deletion suppresses, whereas Rpd3S (*eaf3Δ*) deletion exacerbates, HU sensitivity. **(F)** Epistasis analysis showing that combined deletion of *PHO23* and *RPD3* does not result in further suppression of HU sensitivity in *rad53-K227A* cells, indicating that *PHO23* and *RPD3* function in the same genetic pathway.

To test this model, we assessed how H3K4 methylation and associated reader proteins contribute to fork dynamics using the HU-sensitive *rad53-K227A* checkpoint mutant (Fig. 1A, bottom). We first examined the role of known H3K4me readers, including Pho23, Isw1, Yng1, Set3 (*23, 24*), and the methyltransferase Set1, as well as a histone H3K4A point mutant. Consistent with prior findings, deletion of *SET1* or the H3K4A mutation alleviated HU sensitivity in *rad53-K227A* cells, supporting the idea that H3K4 methylation promotes replication fork reversal upon checkpoint failure (*9*). Strikingly, deletion of *PHO23* also rescued HU sensitivity, whereas deletion of *ISW1*, *SET3*, or *YNG1* had no detectable effect, indicating that Pho23 plays a unique role among H3K4me readers in regulating replication stress (Fig. 1B). In addition to spot dilution assays, we performed growth-curve quantification, showing consistent suppression patterns and quantitatively confirming that *pho23Δ* confers strong rescue of HU sensitivity.

To further dissect the chromatin regulators involved at stalled forks, we examined relevant complexes involved in nucleosome remodeling and histone modification in fork remodeling pathways (Fig. 1C-D). The Rpd3L, recruited through Pho23’s recognition of H3K4me3 (*24*), and the Rpd3S, recruited via Eaf3 binding to H3K36me3 (*13*), were positioned as candidates influencing fork stability during replication stress (Fig. 1D).

To directly test the involvement of H3K4me3 and H3K36me3 reader complexes, we analyzed the HU-sensitivity in *rad53-K227A* background with deletions of *PHO23*, *RXT2*, *CTI6*, *RCO1*, or *EAF3*. Deletion of *PHO23* or *RXT2* strongly suppressed HU sensitivity of *rad53-K227A* cells, indicating that both direct recognition of H3K4me3 by Pho23 and maintenance of Rpd3L structural integrity by Rxt2 are critical for promoting fork instability (*19, 20*). Deletion of *CTI6,* another Rpd3L-specific targeting subunit, resulted in a milder rescue, suggesting a supportive but non-essential role in Rpd3L activity. In contrast, deletion of *RCO1*, a component of Rpd3S, provided only a weak suppression of HU sensitivity, while deletion of *EAF3*, which encodes a chromatin reader anchoring Rpd3S to H3K36me3-marked gene bodies (*25, 26*), exacerbated HU sensitivity in *rad53-K227A* cells (Fig. 1E, top). Growth-curve analyses for *rad53-K227A* strains lacking *PHO23*, *RXT2,* or *EAF3* further corroborated these trends, quantitatively confirming that *Rpd3L* deletion rescues, whereas *Rpd3S* disruption aggravates, HU sensitivity.

To further probe the relationship between H3K4 methylation and Rpd3L function, we combined the H3K4A mutation with deletions of *PHO23* or *RXT2*. Deletion of *PHO23* or *RXT2* alone produced slightly stronger suppression of HU sensitivity than H3K4A alone, and additional suppression was observed when combined with H3K4A (Fig. 1E, bottom). These results suggest that Rpd3L possesses both H3K4 methylation-dependent and methylation-independent activities during replication stress.

To further define the genetic relationship between *PHO23* and *RPD3*, we performed epistasis analysis. Deletion of *PHO23* or *RPD3* alone each suppressed HU sensitivity to a similar extent in *rad53-K227A* cells. Importantly, the double deletion of *PHO23* and *RPD3* only produced slight additive suppression compared to the single mutants (Fig. 1F), indicating that Pho23 and Rpd3 function as a complex to regulate replication fork stability under replication stress. Growth-curve plots for *pho23Δ* and *rpd3Δ* in *rad53-K227A ± H3K4A* backgrounds further support this epistatic relationship, showing overlapping suppression kinetics across HU concentrations. Intriguingly, the H3K4A mutation and *RPD3* deletion exhibit an epistasis relationship, while displaying a minor synergistic effect with *PHO23* deletion in *rad53-K227A* cells (Fig. 1F), suggesting that Rpd3 functions as the primary effector mediating the HU-rescue phenotype.

Together, these findings demonstrate that while H3K4me3 recognition by Pho23 is a major mechanism directing Rpd3L to chromatin during replication stress, Rpd3L also possesses methylation-independent activities that contribute to replication fork destabilization in the absence of functional checkpoint signaling.

### Rpd3L regulates replication fork dynamics through H3K4me-dependent and - independent mechanisms

We next analyzed replication fork dynamics at a site of induced transcription-replication conflict (TRC) using 2D gel electrophoresis. The TRC reporter (*pMET25-3HA-LRE1* adjacent to *ARS305*) was examined under both Met (+) and Met (-) conditions, corresponding respectively to transcription “off” and “on” states (Fig. 2A). Under checkpoint-deficient conditions, 2D gel analysis revealed that wild-type (*RXT2*-H3) cells exhibited strong X-shaped fork reversal signals following hydroxyurea (HU) treatment (Fig. 2B and fig. S1). In the *rxt2Δ*-H3 mutant, fork reversal signals were reduced compared to wild-type, indicating that loss of Rpd3L function partially stabilizes replication forks during replication stress. In the *RXT2*-H3K4A mutant, which eliminates H3K4 methylation, fork reversal signals were even more strongly reduced than in *rxt2Δ*-H3, consistent with the idea that loss of H3K4 methylation exerts a broader or more upstream effect on replication fork stability (*9*). Importantly, in the *rxt2Δ*-H3K4A double mutant, fork reversal signals were further suppressed compared to either single mutant alone (Fig. 2B). Quantification from three independent biological replicates confirmed that fork reversal was progressively reduced across the series from wild-type to *rxt2Δ*-H3 and *RXT2-*H3K4A single mutants, with the double mutant *rxt2Δ*-H3K4A exhibiting the lowest fork reversal signal, indicating additive contributions of Rpd3L function and H3K4 methylation to replication fork destabilization. Consistent with transcriptional regulation, α-HA immunoblotting of 3HA-Lre1 confirmed robust reporter induction under Met (-) but not Met **(+)**, verifying that the fork-reversal phenotype depends on transcription-driven TRC formation.

**Fig. 2.**
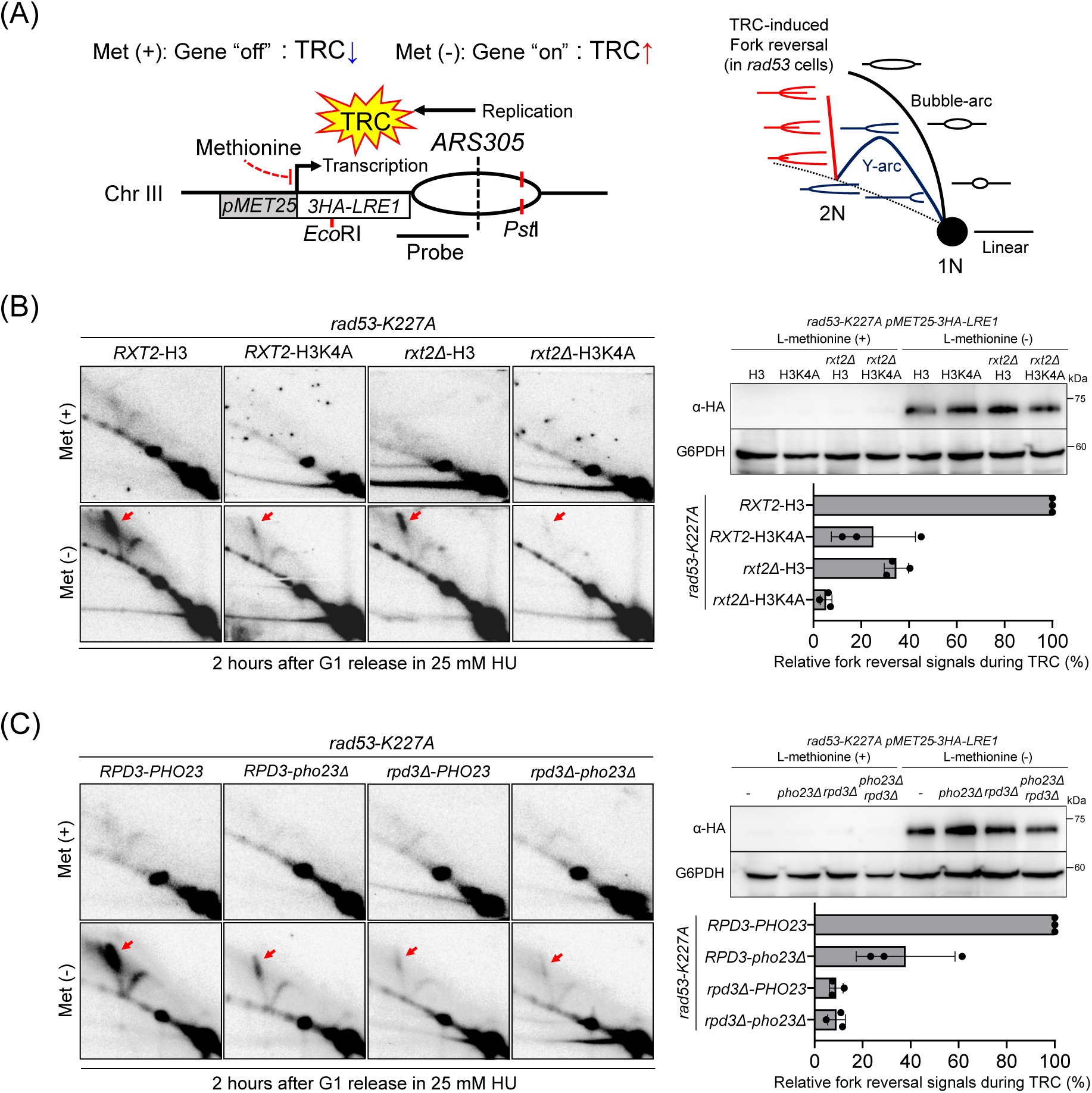
Rpd3L and H3K4 methylation cooperatively regulate replication fork stability during transcription-replication conflicts. All 2D gel analyses were performed in the *rad53-K227A* background. **(A)** Schematic of the experimental system. A head-on transcription–replication conflict (HO-TRC) is induced by activating transcription of the *pMET25-3HA-LRE1* construct integrated near *ARS305* upon methionine depletion (left panel). Replication intermediates are visualized by 2D gel electrophoresis, revealing Y arcs, bubble arcs, and fork reversal structures as indicators of replication stress (right panel). **(B)** 2D gel analysis of replication intermediates in HU-treated cells with the indicated genotypes. Fork reversal signals (red arrows) induced by HO-TRCs upon *LRE1* activation under methionine depletion (upper right panel) are prominent in wild-type (*RXT2*-H3) cells, reduced in *rxt2Δ*-H3 and *RXT2*-H3K4A single mutants, and nearly abolished in the *rxt2Δ*-H3K4A double mutant (left panel). These results indicate cooperative contributions of Rpd3L and H3K4 methylation in restraining fork reversal during transcription–replication conflicts. **(C)** Fork reversal analysis in wild-type (*RPD3*-*PHO23*), *pho23Δ*, *rpd3Δ*, and *rpd3Δ pho23Δ* cells under HU treatment. Fork reversal signals are similarly reduced in *pho23Δ* and *rpd3Δ* single mutants, with no further reduction in the double mutant, indicating that Pho23 and Rpd3 act in the same genetic pathway to resolve replication stress at conflict-prone loci. Fork reversal signals were quantified and are shown as bar charts (upper right of **B** and **C**). Error bars represent mean ± SD.

To further investigate whether Pho23 operates within the same pathway as Rpd3, we compared replication fork reversal phenotypes in checkpoint-deficient (*rad53-K227A*) cells expressing wild-type, *pho23Δ, rpd3Δ*, or *the pho23Δ rpd3Δ* double mutant. Deletion of *PHO23* or *RPD3* individually reduced fork reversal signals relative to wild-type, and notably, the *rpd3Δ pho23Δ* double mutant exhibited a fork reversal phenotype similar to *rpd3Δ* alone (Fig. 2C). Quantitative analysis from three independent experiments showed that deletion of *RPD3* or *PHO23* each significantly reduced fork reversal, and that combining both deletions did not further suppress fork reversal beyond that seen in the *rpd3Δ* single mutant. These results confirm the epistatic relationship between *PHO23 and RPD3*, consistent with the model that Pho23 acts through the Rpd3L complex to modulate replication fork dynamics during transcription-replication conflicts.

Building on previous work showing that H3K4 methylation slows replication forks to prevent transcription-replication conflicts (*9*), these results suggest that Rpd3L regulates replication dynamics through both H3K4 methylation-dependent and -independent mechanisms. Pho23, a core subunit of Rpd3L, binds H3K4me3 via its PHD domain (*19, 20*), supporting recruitment to active chromatin, while Rpd3L also modulates replication fork stability independently of H3K4 methylation. These findings highlight Rpd3L as a chromatin regulator whose functional impact is modulated by checkpoint status during replication stress.

### H3K4 methylation and histone acetylation oppositely regulate Rpd3L-mediated fork stability during replication stress

We next explored whether histone acetylation states directly impact replication fork stability during replication stress. Given that Rpd3L functions as a histone deacetylase complex (HDAC) and Gcn5 is a major histone acetyltransferase (HAT) (*27, 28*), we hypothesized that a balance between HAT-mediated acetylation and HDAC-mediated deacetylation shapes the chromatin environment at TRC-prone loci (Fig. 3A).

**Fig. 3.**
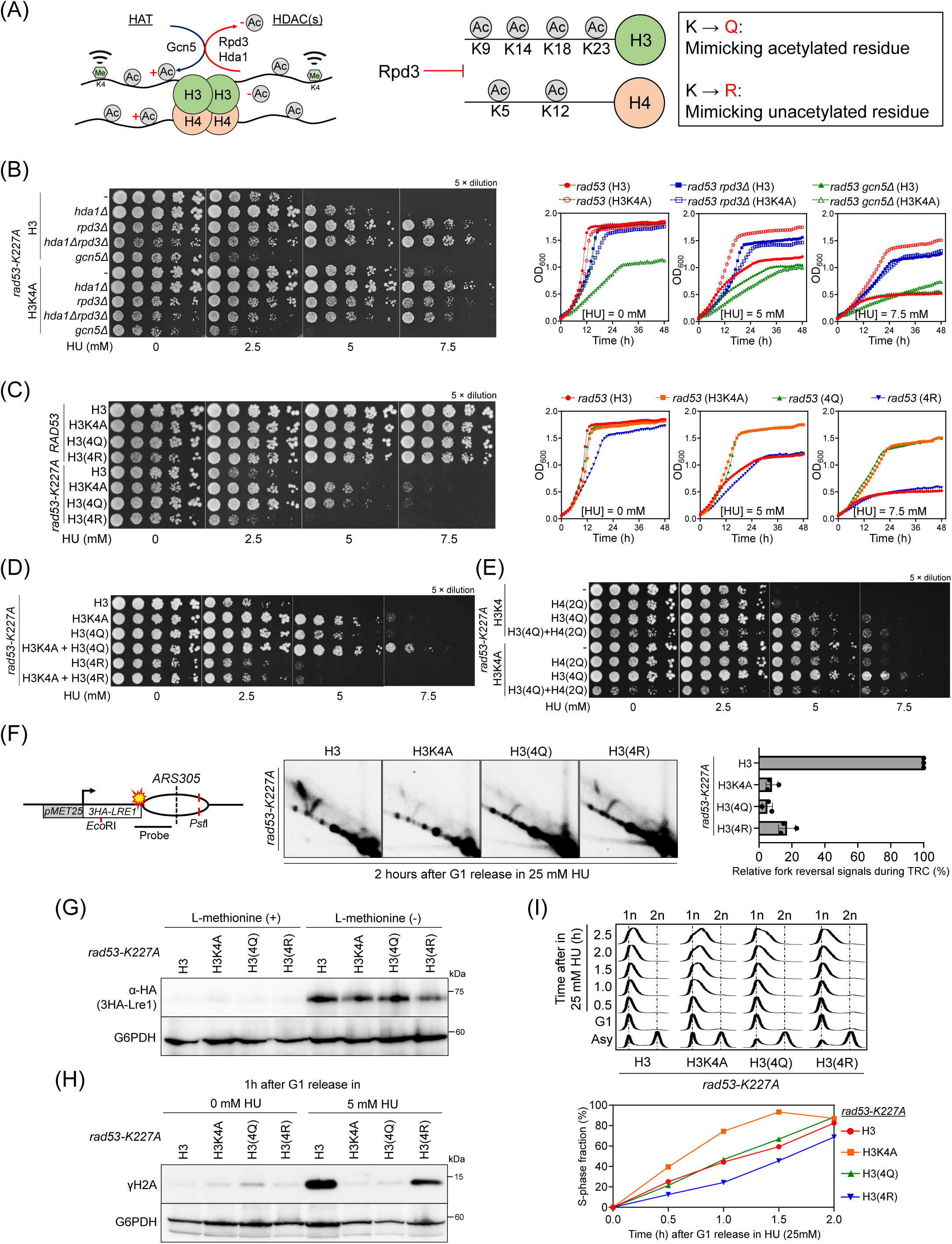
The effects of H3K4 methylation on replication stress sensitivity in *rad53-K227A* mutants are mediated by histone acetylation. (A) Schematic illustration of HAT- and HDAC-mediated regulation of histone H3 and H4 acetylation. Rpd3 functions as an HDAC that deacetylates lysine residues on H3 and H4 tails. Lysine-to-glutamine (K→Q) substitutions mimic acetylated residues, whereas lysine-to-arginine (K→R) substitutions mimic non-acetylated residues. **(B)** Spot dilution assays and growth curves showing HU sensitivity of rad53-K227A cells carrying deletions of *RPD3*, *HDA1*, or *GCN5*, either alone or in combination with H3K4A. Deletion of *RPD3* or *HDA1* suppresses HU sensitivity, whereas deletion of *GCN5* exacerbates HU sensitivity and abolishes H3K4A-mediated suppression. **(C** and **D)** H3(4Q) enhances suppression by H3K4A, whereas H3(4R) diminishes it. **(E)** Genetic combinations of H3K4A, H3(4Q), and H4(2Q) show additive suppression of HU sensitivity, although triple mutants grow more slowly. **(F)** Fork reversal analysis in wild-type (H3), H3K4A, H3(4Q), and H3(4R) cells in the *rad53-K227A* background under HU treatment. Fork reversal signals are reduced in H3K4A and H3(4Q). Quantification is shown (mean ± SD, n = 3). *See also fig. S1A-C*. **(G)** Western blot analysis of Lre1 levels upon transcriptional induction. **(H)** γH2A levels after HU treatment are highest in wild-type cells, reduced in H3K4A and H3(4Q), and remain detectable in H3(4R). **(I)** FACS analysis shows delayed S-phase progression in H3(4R). H3(4Q) and H3(4R) denote histone H3 mutants with lysine-to-glutamine or lysine-to-arginine substitutions, respectively, at K9, K14, K18, and K23.

We found that deletion of *GCN5* markedly exacerbated HU sensitivity in *rad53-K227A* cells, whereas deletion of *RPD3* suppressed the sensitivity, consistent with opposing roles of Gcn5 and Rpd3L in regulating chromatin during replication stress. Deletion of *HDA1*, another histone deacetylase, also conferred suppression but to a lesser extent than *RPD3*, suggesting a more limited or distinct contribution (Fig. 3B). These results support a model in which acetylation protects replication forks under stress, while deacetylation, particularly by Rpd3L, destabilizes forks, in the HU-treated *rad53-K227A* cells.

To further dissect how these chromatin modifiers interact with H3K4 methylation, we combined the H3K4A mutation with deletions of *RPD3*, *HDA1*, or both. Deletion of *HDA1* markedly enhanced the protective effect of H3K4A, suggesting that Hda1 functions largely independently of H3K4 methylation. In contrast, deletion of *RPD3* only mildly increased the suppression conferred by H3K4A, consistent with the idea that Rpd3L is coordinated with H3K4 methylation. Interestingly, the double deletion of *RPD3* and *HDA1* in the H3K4A background resulted in weaker suppression compared to *RPD3* deletion alone, indicating that Rpd3 and Hda1 act through distinct, non-redundant pathways to modulate replication fork dynamics (Fig. 3B, bottom). Conversely, combining H3K4A with *gcn5Δ* diminished the protective effect of H3K4A alone, indicating that Gcn5-mediated acetylation is necessary for the full HU-sensitivity suppression achieved by loss of H3K4 methylation in *rad53-K227A* cells. Growth-curve quantification indicated consistent suppression and enhancement patterns among these genotypes.

To genetically mimic histone acetylation states, we utilized histone H3 point mutants in which lysines 9, 14, 18, and 23 were replaced with either glutamine (H3(4Q)) to mimic acetylation or arginine (H3(4R)) to prevent acetylation (Fig. 3C). To rigorously assess the contribution of each residue, we first examined individual K→Q mutations at H3K9, K14, K18, and K23, either alone or in combination with H3K4A (fig. S2). None of the single mutations conferred detectable suppression of HU sensitivity, even when combined with H3K4A, indicating that acetylation at individual sites is insufficient to promote replication fork stability. These results support the conclusion that the combinatorial acetylation of H3K9, K14, K18, and K23 is required for effective suppression, as further demonstrated by the strong suppression observed with the H3(4Q) mutant (Fig. 3C), a result also supported by quantitative growth-curve analysis showing enhanced suppression kinetics under HU treatment. In contrast, H3(4R) failed to suppress HU sensitivity and significantly diminished the suppression conferred by H3K4A when combined, phenocopying the effect of *gcn5Δ* observed earlier (Fig. 3C-D).

These findings suggest that histone acetylation is required downstream of H3K4A to fully alleviate replication stress in checkpoint-deficient cells, supporting a model in which loss of H3K4 methylation prevents Rpd3L recruitment, while histone acetylation promotes a chromatin environment permissive for replication fork progression.

We next tested whether acetylation of histone H4 contributes to the similarly replication stress response in checkpoint-deficient cells by analyzing combinations of H3(4Q) with H4(2Q), which mimics acetylation at H4 K5 and K12 (Fig. 3E). While H4(2Q) alone conferred only mild suppression of HU sensitivity compared to wild-type H3, its combination with H3(4Q) resulted in slightly enhanced suppression. H3K4A also conferred clear suppression as previously shown, and combining H3K4A with H4(2Q) yielded marginally stronger suppression than H3K4A alone. Importantly, the combination of H3K4A + H3(4Q) resulted in the most robust suppression, underscoring the critical role of histone H3 acetylation in promoting replication fork stability in the absence of H3K4 methylation. The triple-mutant combination of H3K4A + H3(4Q) + H4(2Q) also suppressed HU sensitivity but exhibited slower growth, suggesting that excessive acetylation or the combined removal of multiple chromatin regulatory marks may compromise overall cell fitness despite alleviating replication stress.

To directly assess the impact of histone acetylation states on replication fork stability during TRC, we analyzed strains expressing H3(4Q) and H3(4R) mutants, which mimic acetylated and non-acetylatable states, respectively, at key N-terminal lysines. Both H3K4A and H3(4Q) mutants suppressed HU sensitivity and reduced fork reversal in *rad53-K227A* cells, consistent with a model in which either loss of H3K4 methylation or increased acetylation promotes stability of fragile replication fork (Fig. 3F). Quantification from three independent biological replicates confirmed the reproducibility of fork-reversal suppression. In contrast, H3(4R), which failed to suppress HU sensitivity, displayed only a modest reduction in fork reversal. α-HA immunoblotting of 3HA-Lre1 confirmed equivalent reporter induction under Met (-) conditions across these strains, ruling out transcriptional differences as a cause of the observed effects (Fig. 3G). This modest reduction in fork reversal may reflect slower S phase progression in H3(4R) mutants, which could indirectly reduce TRC formation and associated DNA damage, as indicated by reduced γH2A levels compared to wild-type H3 (Fig. 3H).

Consistent with this, quantitative S-phase completion curves based on FACS analysis confirmed delayed replication progression in H3(4R) compared to wild-type or H3(4Q) cells (Fig. 3I). Notably, the elevated γH2A levels observed in H3(4R) cells, compared to H3K4A and H3(4Q) (Fig. 3H), suggest that significant levels of catastrophic DNA damage may still occur in H3(4R) cells, comparable to wild-type H3 cells.

These results support a model in which H3K4 methylation and Rpd3L-mediated deacetylation functionally interact to regulate chromatin structure and stabilize fragile replication forks resulting from compromised checkpoint signaling. The additive suppression of HU sensitivity observed in H3K4A combined with acetyl-mimic mutations suggests that a chromatin state lacking H3K4 methylation and permissive to acetylation can mitigate transcription-replication conflicts in checkpoint-deficient cells.

### Rpd3L coordinates chromatin acetylation through H3K4 methylation-dependent and independent mechanisms during replication stress

We next investigated how RPD3L regulates chromatin acetylation during replication stress. To eliminate potential interference from checkpoint deficiency, all analyses were performed in *RAD53* wild-type cells, with the corresponding *rad53-K227A* data shown in the fig. S6 for comparison. We first examined global acetylation levels of H3K9/K14 and H3K18, which are primary targets of Gcn5, in cells synchronized in G1 or released into HU for 2 h (Fig. 4A). HU treatment markedly increased both acetylation marks in H3K4A and *pho23Δ* cells, with an even more pronounced increase in *rpd3Δ*. The elevated acetylation in *rpd3Δ* was expected given that Rpd3 is a major histone deacetylase in yeast, but the distinct responses of H3K4A and *pho23Δ* suggest that RPD3L targeting is controlled by both H3K4 methylation-dependent and independent mechanisms. Genome-wide H3K18ac profiles and their quantification are shown in fig. S4 and display trends consistent with those observed for H3K9/K14ac.

**Fig. 4.**
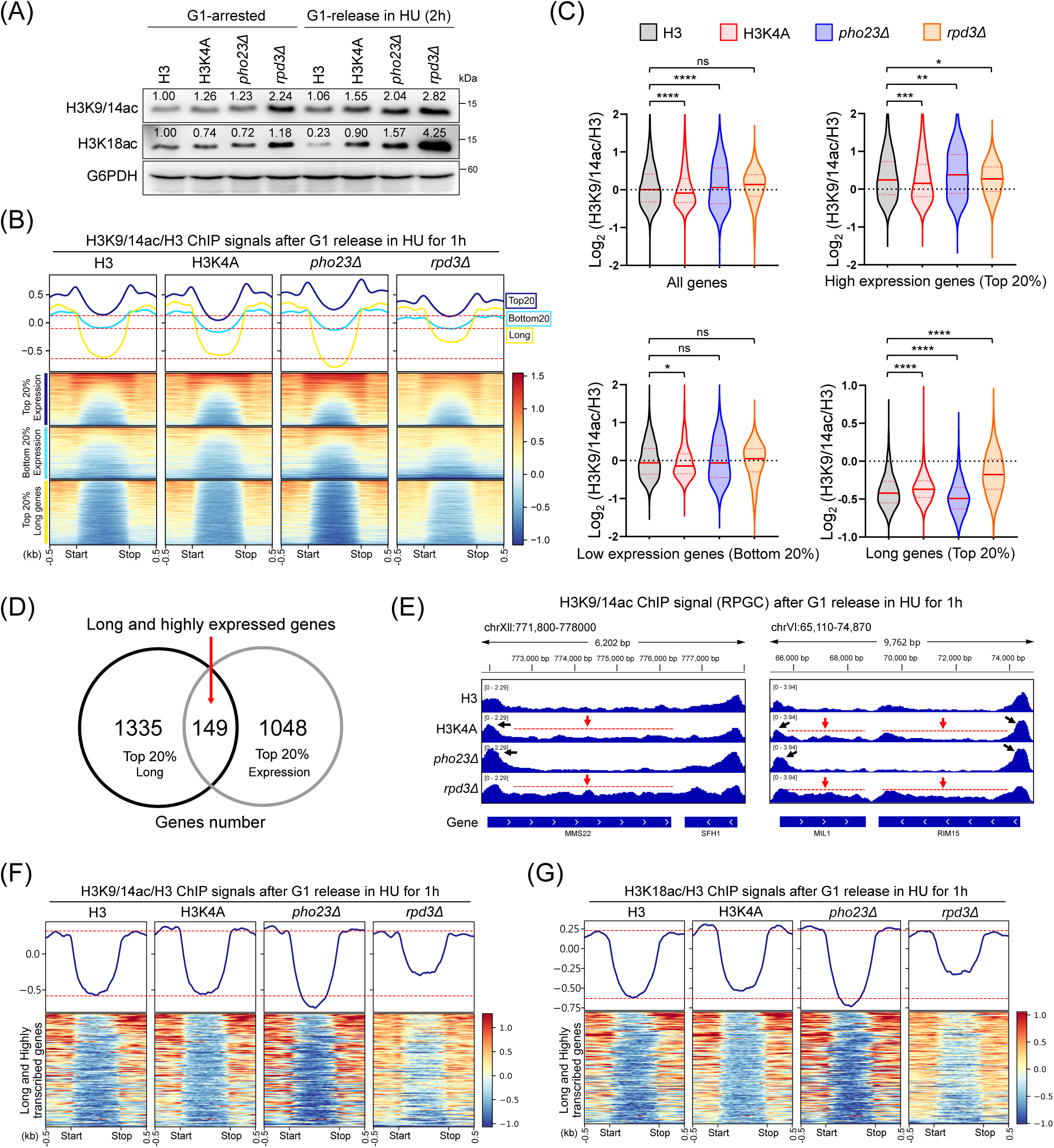
H3K4 methylation and Rpd3L shape H3 acetylation patterns across gene bodies. **(A)** Western blot analysis of H3K9/14ac and H3K18ac in the indicated strains shows elevated overall H3 acetylation levels in H3K4A, *pho23Δ*, and *rpd3Δ* after G1 release in 200 mM HU. **(B)** Metagene profiles and heatmaps showing H3K9/14ac normalized to total H3 ChIP-seq signals after G1 release in HU for 1 h in H3, *H3K4A*, *pho23Δ*, and *rpd3Δ* cells. Signals were calculated across gene bodies (start to stop codon ±500 bp). Heatmaps are shown for genes ranked by expression level (top and bottom 20%) and gene length (top 20% longest genes), revealing distinct H3K9/14ac distribution patterns in H3K4A, *pho23Δ*, and *rpd3Δ* compared with wild-type. **(C)** Violin plots showing genome-wide H3K9/14ac signals normalized to total H3 across gene bodies for all genes, highly expressed genes (top 20%), lowly expressed genes (bottom 20%), and long genes (top 20%). Values represent means from two biological replicates. **(D)** Venn diagram showing overlap between long genes and highly expressed genes. The number of genes in each category and their intersection are indicated. **(E)** Genome browser views of H3K9/14ac signals normalized by RPGC at *MMS22*, *MIL1*, and *RIM15*, representing long and highly expressed genes defined in (D). Red arrows and dashed lines indicate altered acetylation patterns across gene bodies; black arrows mark promoters. **(F** and **G)** Metagene profiles and heatmaps of H3-normalized H3K9/14ac and H3K18ac across long and highly transcribed genes. ChIP-seq signals calculated across gene bodies from the start codon to the stop codon with ±500 bp flanking regions. Statistical significance in (C) was assessed by unpaired Welch’s t test relative to H3 control (ns, \**P* < 0.05, \*\**P* < 0.01, \*\*\**P* < 0.001, \*\*\*\**P* < 0.0001). *See also fig. S4–S6*.

To define how these acetylation changes are distributed across the genome, we performed ChIP-seq for H3K9/K14ac normalized to total H3 (Fig. 4B). H3 ChIP-seq profiles showed a high degree of correlation among wild-type H3, *H3K4A*, *pho23Δ*, and *rpd3Δ* cells following HU treatment (fig. S3), indicating that overall nucleosome occupancy is largely unchanged. We analyzed the entire coding sequences (CDS) with flanking 0.5 kb upstream and downstream regions, reasoning that transcription-replication conflicts (TRCs) are more likely to occur within gene bodies where replication forks encounter elongating RNA polymerase. The average global levels of H3K9/K14ac exhibited a significant reduction in *H3K4A* cells and an increase in *pho23Δ* cells, and surprisingly showed no significant change in *rpd3Δ* cells despite the dramatic differences observed by western blotting (Fig. 4A). Equivalent genome-wide analyses in the *rad53-K227A* background are shown in fig. S6 and reveal patterns comparable to those observed in *RAD53* wild-type cells.

To assess whether these changes are linked to transcriptional activity, we stratified genes into the top and bottom 20% by expression level, as well as the top 20% longest genes, a class of loci most susceptible to TRCs (*3*) (fig. S3 and Materials and Methods). In H3K4A cells, acetylation decreased in highly expressed genes but increased in long genes, whereas in *pho23Δ*, acetylation increased in highly expressed genes but decreased in long genes (Fig. 4C). These reciprocal patterns indicate that the RPD3L-Pho23 interaction with H3K4 methylation limits RPD3L activity to H3K4me-enriched regions of highly transcribed genes, while its absence redistributes RPD3L function to long transcription units. In *rpd3Δ*, acetylation increased moderately at highly expressed genes but rose dramatically across long genes, demonstrating that RPD3L also exerts H3K4me-independent control of chromatin acetylation along extended gene bodies. Consistent stratified analyses for H3K18ac are presented in fig. S5.

We next focused on the 149 genes that are both long and highly expressed (Fig. 4D), a class of loci most susceptible to TRCs as shown in mammalian systems (*29*). At representative genes (*MMS22*, *RIM15*), HU-induced acetylation increases were restricted to promoters in *pho23Δ*, extended into coding regions in H3K4A, and expanded across both regions in *rpd3Δ* (Fig. 4E). Metagene analyses of all 149 long, highly expressed genes confirmed this trend (Fig. 4F), and similar patterns observed for H3K18ac (Fig. 4G) further support that RPD3L modulates acetylation through multiple targets of Gcn5. Together, these data demonstrate that RPD3L coordinates chromatin acetylation through dual mechanisms: an H3K4 methylation-dependent pathway mediated by Pho23 at active promoters and an H3K4 methylation-independent pathway acting across long genes where transcription-replication conflicts are most frequent.

### H3K4 methylation promotes Rpd3L recruitment to highly transcribed genes through Pho23

To examine how Rpd3L is targeted to chromatin, we analyzed the genome-wide occupancy of its H3K4me-binding subunit Pho23 by ChIP-seq. Because tagging may interfere with protein function, we first evaluated whether different epitope tags on Pho23 or Rpd3 affect their biological activity in the *rad53-K227A* background, where HU sensitivity provides a sensitive readout for Rpd3L function (Fig. 5A). The growth assays showed that only Pho23-9×Myc did not alter the HU sensitivity of *rad53-K227A* cells, whereas other tagged versions of Pho23 or Rpd3 compromised function to varying degrees. Therefore, all subsequent ChIP-seq analyses were performed using Pho23-9×Myc as a representative and functionally intact reporter for Rpd3L occupancy (illustrated in the schematic, Fig. 5A, right panel).

**Fig. 5.**
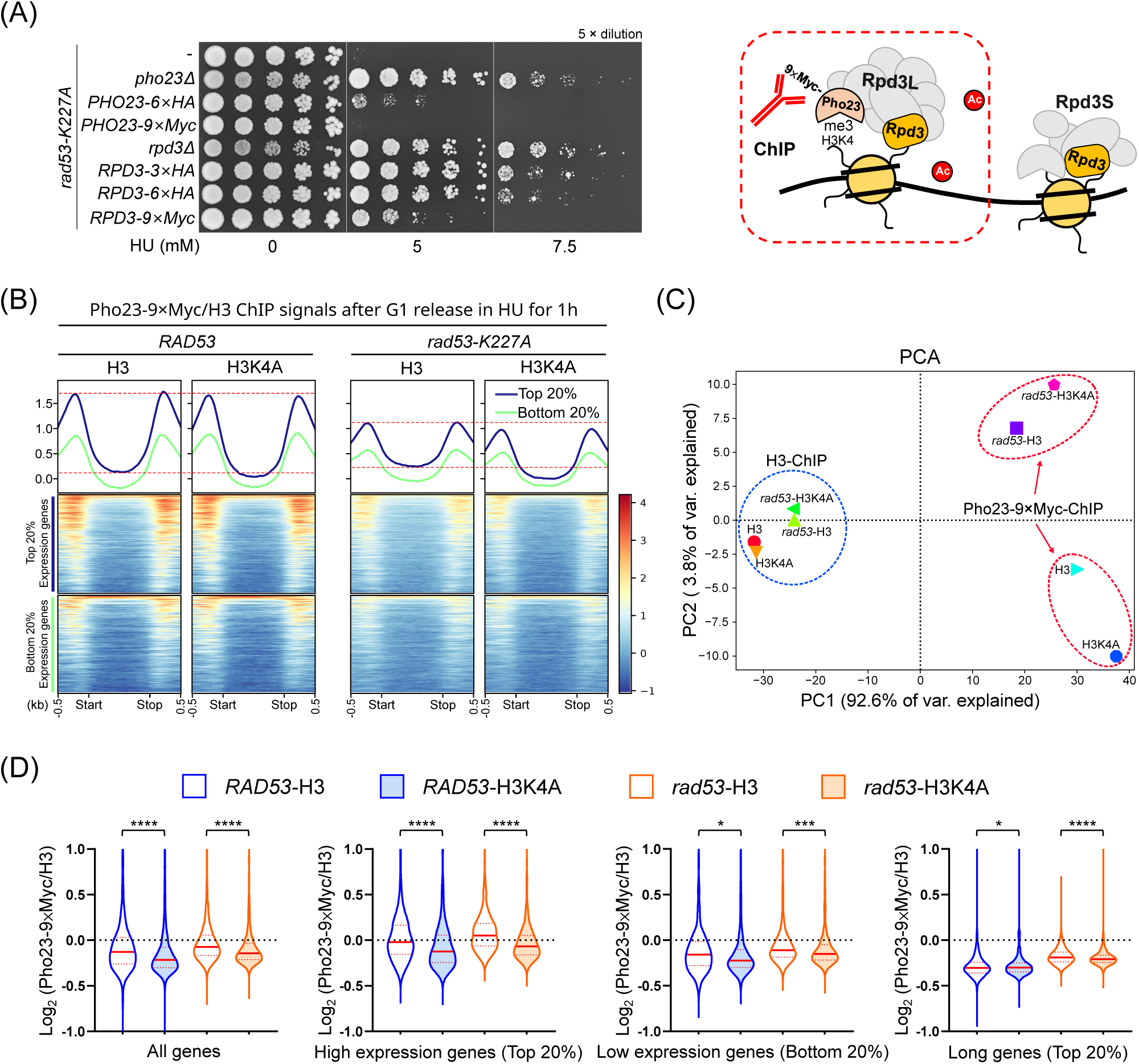
H3K4 methylation facilitates Rpd3L association with chromatin. **(A)** Spot dilution assays showing HU sensitivity of *rad53-K227A* cells carrying the indicated deletions or C-terminally epitope-tagged *PHO23* and *RPD3* alleles. A schematic illustrating the use of Pho23-9×Myc as a reporter for Rpd3L chromatin association is shown on the right. **(B)** Metagene profiles and heatmaps showing Pho23-9×Myc ChIP-seq signals normalized to total H3 after G1 release in HU treatment in *RAD53* (200 mM) and *rad53-K227A* (25 mM) cells carrying H3 or H3K4A. Signals were calculated across gene bodies (start to stop codon ±500 bp). Genes are grouped by expression level (top and bottom 20%). **(C)** Principal component analysis (PCA) of RPGC-normalized ChIP-seq signals from Pho23-9×Myc ChIP and H3 ChIP datasets, showing separation by Rad53 status and H3K4 methylation state. H3 ChIP datasets cluster closely across strains, indicating comparable nucleosome occupancy. **(D)** Violin plots showing genome-wide H3-normalized Pho23-9×Myc ChIP-seq signals across whole gene bodies in *RAD53* and *rad53-K227A* cells carrying H3 or H3K4A. Data are shown for all genes, highly expressed genes (top 20%), lowly expressed genes (bottom 20%), and long genes (top 20%). Values represent means from two biological replicates. Statistical significance was assessed by unpaired Welch’s *t* test relative to the corresponding H3 control (ns, * *P* < 0.05; ** *P* < 0.01; *** *P* < 0.001; **** *P* < 0.0001).

Pho23-9×Myc ChIP-seq normalized to total H3 revealed that Pho23 occupancy was reduced in all gene categories in H3K4A cells compared to H3 wild-type cells following HU treatment, but the most pronounced loss occurred at highly expressed genes in both *RAD53* wild type and *rad53-K227A* cells (Fig. 5B, D). These results are consistent with the observation that H3K9/K14 acetylation levels are most strongly affected in highly transcribed genes in *pho23Δ* cells (see Fig. 4B), suggesting that RPD3L is recruited by the Pho23-H3K4me interaction to highly transcribed genes.

Principal-component analysis (PCA) showed that H3 ChIP-seq datasets from all strains were highly similar, whereas Pho23-9×Myc ChIP profiles separated along the first component according to genotype (Fig. 5C). This indicates that the observed differences in Pho23 occupancy reflect specific regulatory changes rather than alterations in chromatin accessibility. Collectively, these results demonstrate that Rpd3L targeting is mediated by H3K4 methylation through Pho23, particularly at highly expressed genes, supporting the conclusion that Rpd3L coordinates chromatin state via both H3K4 methylation-dependent and independent mechanisms during replication stress.

### H3K4 methylation and Rpd3L restrain replication dynamics and suppress transcription-associated mutagenesis

We next evaluated whether H3K4 methylation and Rpd3L regulate replication dynamics and genome stability beyond the checkpoint-defective conditions. All assays were performed in an intact *RAD53* background to assess their physiological functions under replication stress. We first assessed replication fork progression by 2D gel electrophoresis and S-phase progression by flow cytometry in cells expressing wild-type H3, H3K4A, H3(4Q), or lacking *PHO23* (Fig. 6A). In wild-type H3 cells, bubble arc signals and moderate Y arc signals were detected, reflecting regulated origin firing and fork progression. In contrast, H3K4A, H3(4Q), and *pho23Δ* strains displayed diminished bubble arc intensity and stronger Y arc signals, indicative of faster replication fork movement. Quantification of S-phase progression confirmed that these mutant cells completed a higher proportion of DNA synthesis after 1 h of HU exposure compared with wild-type (Fig. 6B). Together, these findings suggest that loss of H3K4 methylation or Pho23 function accelerates replication dynamics under replication stress.

**Fig. 6.**
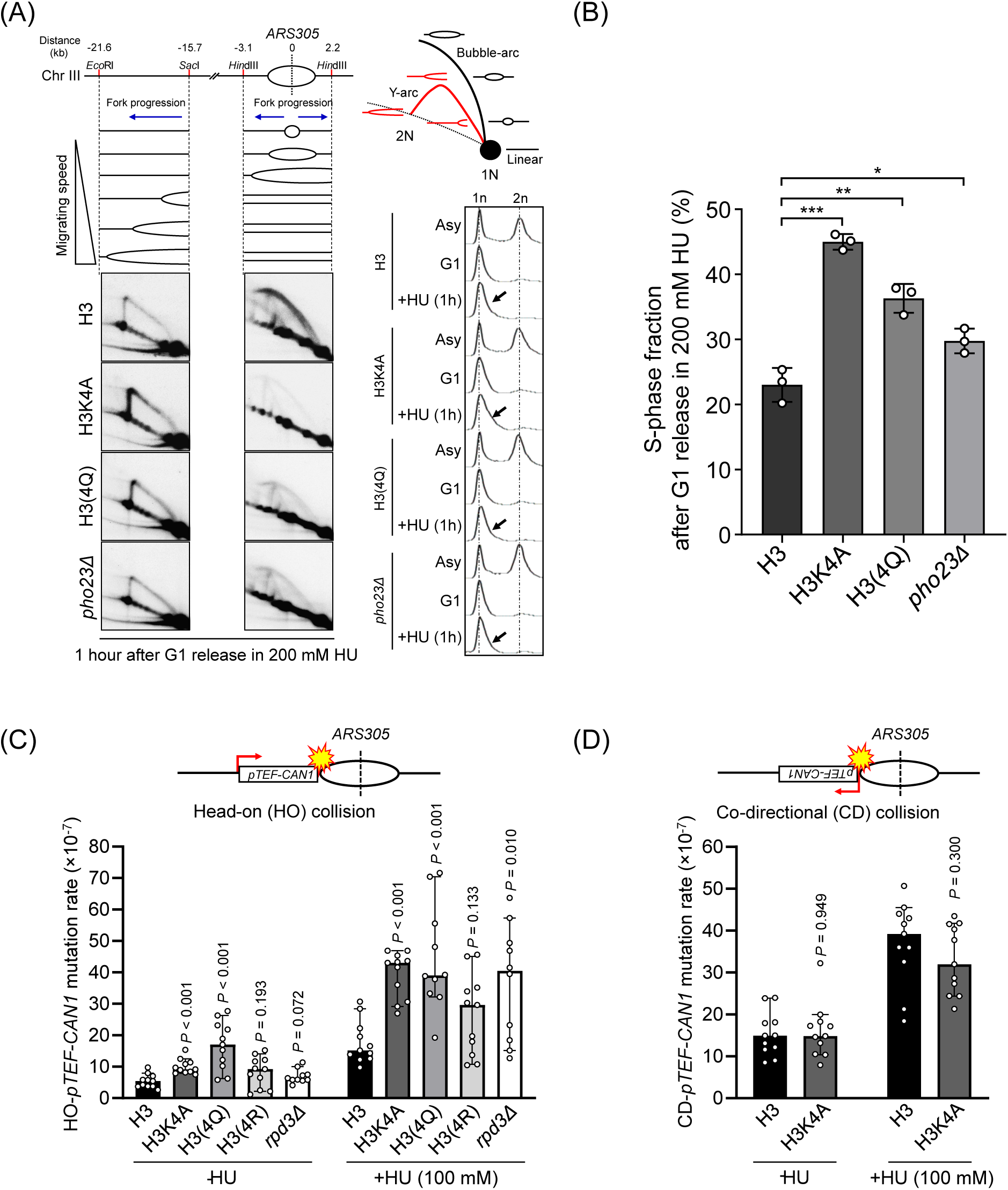
Rpd3L function restrains replication fork progression and maintains genome stability under replication stress. **(A)** 2D gel analysis of replication intermediates at regions originating from *ARS305* in the indicated strains (H3, H3K4A, H3(4Q), and *pho23Δ*) following G1 synchronization and release into 200 mM HU for 1 h. Schematics indicate the analyzed restriction fragments, replication fork direction, and the positions of bubble arc and Y arc species. Compared with H3 cells, bubble arc signals are reduced and Y arc signals are elevated in H3K4A, H3(4Q), and *pho23Δ* strains, consistent with accelerated replication fork progression. Representative FACS profiles are shown on the right, indicating faster S-phase progression in the mutant strains (black arrows). **(B)** Quantification of the S-phase fraction after G1 release in 200 mM HU in the indicated strains. This analysis extends the 2D gel experiment shown in panel (A), assessing S-phase progression by flow cytometry 1 h after G1 release in 200 mM HU. Data represent mean ± SD from three independent biological replicates (n = 3). Statistical significance was assessed by Student’s *t* test, with comparisons performed relative to the H3 (wild-type) control. **(C** and **D)** Schematic illustrations of head-on (HO) and co-directional (CD) transcription–replication conflict (TRC) configurations generated by integrating a *pTEF***-**driven *CAN1* reporter adjacent to *ARS305*. Mutation rates of the HO-*pTEF*-*CAN1* (C) and CD-*pTEF*-*CAN1* (D) reporters were measured in the indicated strains grown in the absence (-HU) or presence (+HU, 100 mM) of hydroxyurea. Data are presented as the median with 95% confidence intervals from 11 independent samples (n = 11). Statistical significance relative to the H3 control was determined using the Mann-Whitney *U* test, and exact *P* values are indicated.

To determine whether this altered replication progression is associated with increased genome instability, we analyzed mutation frequencies using a *pTEF*-driven *CAN1* reporter inserted adjacent to *ARS305*. The reporter was positioned either in a head-on or co-directional orientation relative to replication forks (Fig. 6C-D). Using this system, we measured the spontaneous mutation frequency. In the absence of HU, H3(4Q) mutants exhibited significantly elevated mutation rates, suggesting that mimicry of elevated H3 acetylation compromises genome integrity at highly transcribed loci. Under HU treatment, all three mutants-H3K4A, H3(4Q), and *rpd3Δ*-displayed higher mutation frequencies than wild-type, indicating that both H3K4 methylation and Rpd3L contribute to suppressing transcription-associated mutagenesis during replication stress. In contrast, no significant differences were detected in the co-directional orientation (Fig. 6D), indicating that replication-induced genome instability primarily arises from head-on transcription-replication conflicts.

## Discussion

Epigenetic regulation of chromatin structure plays a critical role in coordinating transcription and replication programs. In this study, we uncover a chromatin-based mechanism in which the Rpd3L histone deacetylase complex modulates replication fork dynamics and preserves genome stability through both H3K4 methylation-dependent and H3K4 methylation-independent modes of chromatin engagement. We propose a working model (Fig. 7) in which Rpd3L activity dynamically modulates chromatin state at actively transcribed regions under replication stress. In wild-type cells (Fig. 7A), H3K4 methylation promotes Pho23-mediated recruitment of Rpd3L to these regions, establishing a restrictive chromatin state that slows replication forks and mitigates transcription-replication conflicts (TRCs). In parallel, Rpd3L can also engage chromatin independently of H3K4 methylation, particularly under stress, contributing further to chromatin restriction and fork slowing. Conversely, loss of H3K4 methylation (through *SET1* deletion or H3K4A mutation; Fig. 7B) diminishes Rpd3L recruitment and allows Gcn5-mediated histone acetylation to dominate, promoting a permissive chromatin state that accelerates replication fork progression but increases the risk of TRCs, replication errors, and transcription-associated mutagenesis. These findings demonstrate that chromatin state modulation through coordinated histone methylation, deacetylation, and acetylation dynamically controls replication fork progression and transcription-replication conflict resolution. Together, our results define an integrated chromatin-based mechanism that protects genome stability and suppresses transcription-associated mutagenesis under replication stress by balancing restrictive and permissive chromatin states at actively transcribed regions.

**Fig. 7.**
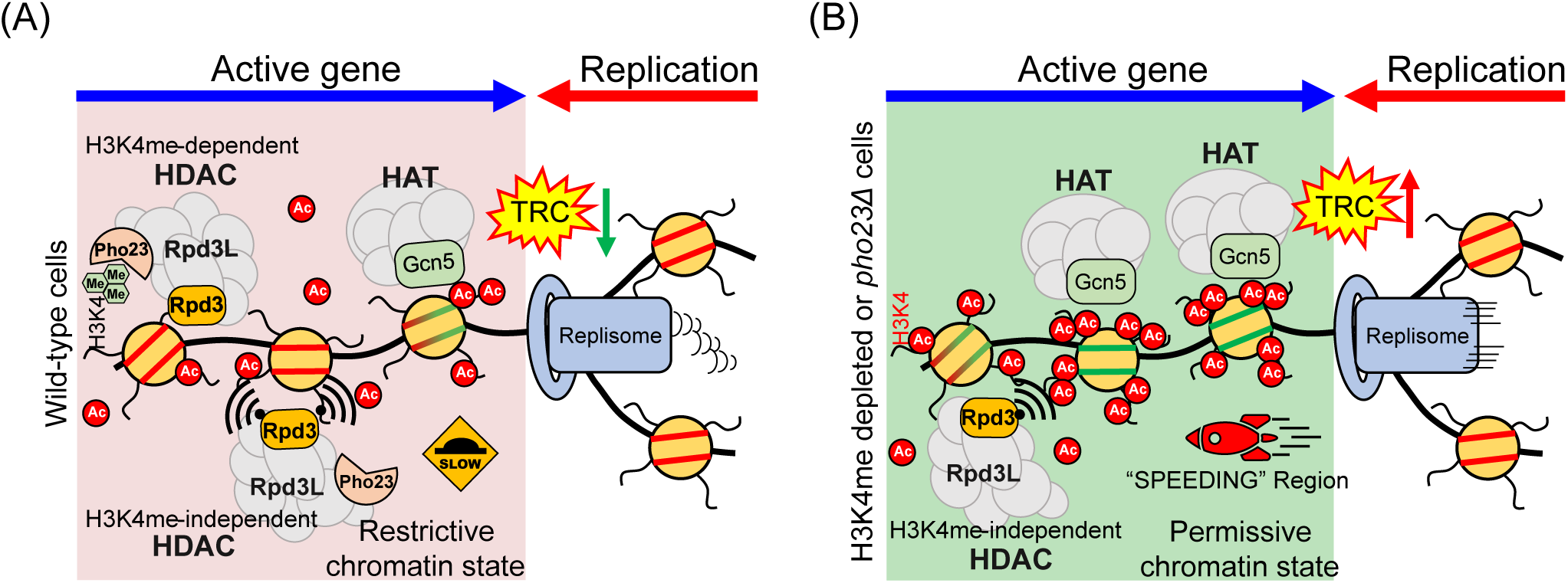
Working model for Rpd3L-mediated regulation of replication fork progression at transcription-replication conflicts. **(A)** In wild-type cells, H3K4 methylation at actively transcribed regions promotes Pho23-mediated recruitment of the Rpd3L complex, enabling H3K4 methylation-dependent histone deacetylation across gene bodies. This activity counterbalances HAT-driven acetylation and establishes a relatively restrictive chromatin state that slows replication fork progression and mitigates TRCs. In parallel, Rpd3L can also associate with chromatin through H3K4 methylation-independent mechanisms, providing additional basalHDAC activity that contributes to fork restraint under stress conditions. **(B)** In the absence of H3K4 methylation or Pho23, H3K4 methylation-dependent recruitment of Rpd3L is compromised, allowing Gcn5-mediated histone acetylation to dominate. This shift toward a permissive chromatin state facilitates accelerated replication fork progression through actively transcribed regions, increasing the likelihood of unresolved TRCs and transcription-associated genome instability.

This study provides new insight into how chromatin structure modulates replication dynamics and genome stability. While H3K4 methylation is traditionally viewed as a transcription-associated mark (*30*), our findings demonstrate that it also contributes to replication fork control by regulating chromatin acetylation and HDAC recruitment. Importantly, we identify Rpd3L but not Rpd3S as the functionally relevant complex in this context, reinforcing the principle that chromatin-modifying complexes with shared catalytic subunits can perform distinct biological roles based on their scaffold composition and recruitment pathways (*31*). Structural studies show that Pho23 binds H3K4me3 through its PHD domain (*19, 20*), consistent with our ChIP-seq evidence that Pho23 occupancy is reduced in H3K4A mutants. This chromatin axis also overlaps with genome maintenance functions of Rpd3L described in earlier genetic studies: loss of Rpd3L improves checkpoint-deficient cell survival (*16*) and restores DNA damage response in checkpoint-deficient mutants (*17*). Furthermore, Rpd3L cooperates with Hda1 to promote sister chromatid cohesion and repair broken forks via cohesion establishment (*16*). Together with recent studies emphasizing the role of chromatin features in limiting transcription-replication interference (*4, 32*), our findings position Rpd3L as a central mediator linking histone methylation to chromatin state adaptation under replication stress.

The dual roles of Rpd3L in transcriptional repression and replication fork regulation appear to be evolutionarily conserved. In mammalian cells, ING2, a PHD finger protein and component of the Sin3A-HDAC1/2 complex, directly binds H3K4me3 through a structurally conserved reader domain (*22*) and facilitates gene repression via histone deacetylation (*21*). Beyond transcription, ING2 also promotes replication fork progression by enhancing PCNA chromatin loading, and its loss leads to fork slowing, S-phase defects, and genome instability (*33*). Further supporting a conserved chromatin-mediated mechanism of genome maintenance, several studies have shown that human SETD1A-dependent H3K4 methylation protects stalled replication forks from nucleolytic degradation by promoting FANCD2-mediated histone chaperone activity and RAD51 loading (*34*). Moreover, mutations in MLL2/KMT2D result in transcriptional stress and genome instability by impairing H3K4me3 near transcription start sites and enhancing TRC formation (*35*). Another recent study showed that H3K4me1 regulated by KMT2C/D modulates replication origin activity, highlighting broader roles for H3K4 methylation in S-phase control (*36*). These findings across species point to a conserved strategy whereby H3K4 methylation contributes to fork protection, transcription-replication coordination, and genome stability through chromatin-based regulation.

While this study establishes a mechanistic link between H3K4 methylation, Rpd3L, and replication fork regulation, several important questions remain. First, the genome-wide specificity of Rpd3L targeting under stress is not fully defined. Although Pho23-H3K4me3 interaction contributes to its recruitment, our data suggest additional H3K4me-independent mechanisms-possibly involving stress-induced chromatin changes-that require further characterization. Second, the precise genomic loci where this pathway mitigates TRCs remain to be identified. Genome-wide mapping of TRC-prone regions or transcription-replication interference sites could provide further mechanistic insight. Third, while our work focuses on budding yeast, components of this regulatory axis-H3K4me3, HDAC1/2, and TRC resolution factors-are highly conserved. Future studies should explore whether analogous mechanisms operate in mammalian systems, where TRC management is particularly critical in highly proliferative or transcriptionally active tissues, such as during development (*37*) or in cancer (*38, 39*).

### Materials and Methods Yeast strain construction

All yeast strains carrying the indicated H3 mutations were derived from the MSY421 background (39), a strain in which all genomic copies of the H3-H4 genes are deleted and viability is maintained by extrachromosomal expression of H3-H4 genes from the plasmid pMS329 [*CEN4 ARS1 URA3 HHT1 HHF1*]. All H3 and H4 mutant alleles were generated by site-directed mutagenesis using the plasmid pMS337 [*CEN4 ARS1 LEU2 HHT1 HHF1*]. To generate yeast strains harboring the indicated H3 and H4 mutations, the H3-H4 genes carried on pMS329 were replaced with either wild-type or mutant H3-H4 alleles from pMS337 using standard plasmid shuffling procedures. For ChIP-seq analyses of the RPD3L complex, a 9×Myc epitope tag was fused to the C-terminus of the endogenous *PHO23* gene by homologous recombination using a PCR fragment amplified from pYM20 (*40*). For mutation assay experiments, a *pTEF*-driven *CAN1* cassette flanked by homology arms was amplified from a strain described in our previous study (*9*) and integrated to replace the *URA3* marker that had been pre-integrated adjacent to *ARS305* in a *can1* deletion background. Transformants were selected on 5-fluoroorotic acid (5-FOA)-containing plates. This strategy generated marker-free head-on (HO) and co-directional (CD) *pTEF-CAN1* reporter strains. Detailed descriptions of all yeast strains, plasmids, and primers used for site-directed mutagenesis and strain construction are provided in Tables S1-3.

### Yeast spot dilution assay

Overnight liquid cultures grown in YPD medium were diluted into fresh YPD to an initial OD₆₀₀ of 0.2 and allowed to grow for 5 h at 30 °C to reach log phase. Cells were then collected, resuspended to a final concentration of 2.5 × 10⁶ cells/mL, and subjected to fivefold serial dilutions in distilled water. An aliquot of 2.5 μL from each dilution was spotted onto YPD agar plates supplemented with hydroxyurea (HU) at the indicated concentrations. Plates were incubated at 30 °C for 3 days, and cell growth was recorded.

### Yeast growth curve analysis

Yeast strains were grown overnight in YPD medium at 30 °C with shaking at 200 rpm. Saturated cultures were diluted into fresh YPD medium to an initial OD_600_ of 0.1. A 2× hydroxyurea (HU) stock medium was prepared by dissolving HU in YPD to a final concentration of 400 mM. Lower HU concentrations (200, 15, and 10 mM) were obtained by serial dilution with YPD. For growth curve assays, cultures were dispensed into 96-well microplates by mixing equal volumes (100 μL each) of cell suspension and HU-containing medium, resulting in a final OD_600_ of 0.05 and HU concentrations at half of the stock medium (e.g., a 400 mM stock yielding 200 mM final HU). Cell growth was monitored using an absorbance microplate reader (BMG SPECTROstar Nano) at 30 °C with double-orbital shaking at 400 rpm. Optical density at 600 nm was measured every hour for 48 hours. Each condition was measured in technical triplicate.

### DNA Two-dimensional (2D) gel electrophoresis

All yeast strains used for 2D gel analysis were *BAR1*-deleted. Log-phase cells (OD_600_ ≈ 0.5) were synchronized in G1 phase by treatment with α-factor (100 ng/mL) for 2.5 hours in the appropriate medium. Cells were washed and released into fresh medium containing the indicated concentrations of hydroxyurea (HU) at 30 °C. Approximately 3 × 10^10^ cells were harvested at the indicated time points, and replication intermediates were immediately stabilized by the addition of 0.1% (w/v) sodium azide. Cells were resuspended in 5 mL ice-cold water, transferred to six-well plates, and incubated with 300 μL trioxsalen solution (200 μg/mL in 100% ethanol) for 10 minutes, followed by irradiation with UVA light (365 nm) for 10 minutes on ice. This cross-linking procedure was repeated four times. Cells were then spheroplasted in 5 mL yeast lysis buffer (1 M sorbitol, 100 mM EDTA, 14 mM β-mercaptoethanol, and 5 mg Zymolyase®-100T) at 37 °C for 1 hour. Spheroplasts were collected and resuspended in 4 mL digestion buffer (800 mM guanidine HCl, 30 mM Tris-HCl [pH 8.0], 30 mM EDTA, 5% Tween-20, and 0.5% Triton X-100) supplemented with RNase A (100 μL of 10 mg/mL), followed by incubation at 37 °C for 1 hour. Protein digestion was performed by adding 200 μL proteinase K (20 mg/mL) and incubating at 50 °C for 3 hours, followed by incubation at 30 °C overnight. Genomic DNA was purified using Genomic-tip 100/G columns (QIAGEN, Cat. No. 10243). Briefly, supernatants were diluted with an equal volume of equilibration buffer (750 mM NaCl, 50 mM MOPS [pH 7.0], 15% isopropanol, and 0.15% Triton X-100) and loaded onto columns pre-equilibrated with 4 mL equilibration buffer. Columns were washed twice with 7.5 mL wash buffer (1.0 M NaCl, 50 mM MOPS [pH 7.0], and 15% isopropanol), and genomic DNA was eluted with 5 mL elution buffer (1.25 M NaCl, 50 mM Tris-HCl [pH 8.5], and 15% isopropanol) at 50 °C, followed by isopropanol precipitation. Purified DNA (8 μg) was digested with the indicated restriction enzyme(s), isopropanol-precipitated, and resuspended in 20 μL TE buffer. DNA samples were resolved by first-dimension electrophoresis on a 0.35% agarose gel at 1.5 V/cm for 19 hours, followed by second-dimension electrophoresis on a 1% agarose gel at 6 V/cm for 5 hours in TBE buffer, using a Maxi Horizontal Gel Electrophoresis System (Major Science, ME20-25). DNA was transferred to membranes by standard Southern blotting procedures. PCR-amplified DNA fragments corresponding to the indicated genomic regions were used as probes; primer sequences are listed in Table S4. Hybridized membranes were exposed to Storage Phosphor Screens (GE Healthcare, BAS-IP MS 2040 E) for 5 days, and signals were detected using a Typhoon™ FLA 7000 Phosphorimager (GE Healthcare) at 1000 PMT voltage. Fork reversal signal intensities were quantified using ImageJ. Data from three biological replicates were normalized to the indicated strain and visualized as bar charts using GraphPad Prism. The quantification method and biological replicate data are shown in fig. S1.

### Western blot analysis

Total cell lysates were prepared using the trichloroacetic acid (TCA) protein extraction method. Cells were resuspended in TCA buffer (1.85 M NaOH and 7.4% β-mercaptoethanol) and incubated for 10 minutes, followed by the addition of an equal volume of 20% trichloroacetic acid to precipitate proteins. Precipitated lysates were collected and resuspended in 0.1% NaOH. Protein samples were resolved by SDS-PAGE and transferred onto PVDF membranes (Immobilon®-P, Millipore). Membranes were incubated with the indicated primary antibodies. The following antibodies were used in this study (dilution, source, and catalog number): anti-HA (1:5,000, Roche, 11867423001), anti-H2A S129 phosphorylation (1:2,000, Abcam, ab15083), anti-H3K9/14ac (1:5,000, Merck Millipore, 06-599), anti-H3K18ac (1:3,000, Abcam, ab1191), and anti-G6PDH (1:10,000, Sigma, A9521). Immunoreactive signals were detected using an enhanced chemiluminescence (ECL) detection reagent (Immobilon™, Millipore) and visualized with a UVP BioSpectrum® imaging system.

### Flow cytometry analysis of cell cycle progression

Cells carrying the *bar1Δ* mutation were synchronized in G1 phase at the onset of log phase by treatment with α-factor (100 ng/mL) for 2.5 hours and subsequently released into the indicated experimental conditions. Cells were collected at the specified time points and fixed in 70% ethanol. Fixed cells were treated with RNase A (1 mg/mL) in 50 mM Tris-HCl (pH 8.0) at 37 °C to remove RNA. Cells were washed and subsequently treated with proteinase K (2 μL of >800 units/mL) in 100 μL of 30 mM Tris-HCl. DNA was stained with SYBR™ Green I (1:3,000 dilution; Invitrogen, S7563) in 50 mM Tris**-**HCl (pH 8.0) by incubation overnight at 4 °C. Samples were analyzed using a flow cytometer (Thermo Fisher Scientific, Attune™ NxT). For each time point, 20,000 gated events were collected and analyzed using FlowJo software. The proportion of cells in S phase was determined using the Dean-Jett-Fox model. Data from three independent biological replicates were visualized as bar charts using GraphPad Prism.

### Chromatin immunoprecipitation sequencing (ChIP-seq)

G1-arrested *RAD53* cells were treated with 200 mM hydroxyurea (HU) for 1 hour, whereas *rad53* cells were treated with 25 mM HU for 1 hour, prior to harvesting and cross-linking for chromatin immunoprecipitation. Cells were fixed with formaldehyde fixation solution (5 mM HEPES, pH 7.6; 0.1 mM EDTA; 10 mM NaCl; and 1.1% formaldehyde) for 20 minutes at room temperature and subsequently quenched with 125 mM glycine for 5 minutes. Cell pellets were washed and resuspended in 10 mL of ice-cold TBS (20 mM Tris, pH 7.6, and 150 mM NaCl). Cells were disrupted using a homogenizer (FastPrep®-24 5G) with the built-in program optimized for *Saccharomyces cerevisiae*. Chromatin was sheared by sonication using a Diagenode Bioruptor® Pico sonicator to generate DNA fragments with an average size of approximately 200–500 bp in 700 μL of FA buffer (50 mM HEPES-KOH, pH 7.6; 150 mM NaCl; 1 mM EDTA; 1% Triton X-100; 0.1% sodium deoxycholate; and 1× protease inhibitor cocktail). Equal volumes of chromatin were subjected to immunoprecipitation. For H3K9/14ac ChIP, 4 μL of anti-H3K9/14ac antibody (Merck Millipore, 06-599) was conjugated to 20 μL of Protein G magnetic beads (Cytiva Protein G Mag Sepharose™). For H3K18ac ChIP, 5 μL of anti-H3K18ac antibody (Abcam, ab1191) was conjugated to 20 μL of Protein G magnetic beads. For total H3 ChIP, 4 μL of anti-H3 antibody (Abcam, ab1791) was conjugated to 20 μL of Protein G magnetic beads. For Myc-tagged protein ChIP, 4 μL of anti-Myc antibody (Merck Millipore, 05-419) was conjugated to 20 μL of Protein A magnetic beads (Cytiva Protein A Mag Sepharose™). Antibody-conjugated beads were resuspended in 30 μL of PBS-T for immunoprecipitation assays. For each immunoprecipitation, 200 μL of sonicated chromatin was incubated with antibody-conjugated beads at 4 °C overnight. Chromatin-bound beads were washed three times with FA buffer, twice with FA-HS buffer (FA buffer containing 500 mM NaCl), and once with RIPA buffer (10 mM Tris, pH 8.0; 250 mM LiCl; 0.5% NP-40; 0.5% sodium deoxycholate; 1 mM EDTA; and 1× protease inhibitor cocktail). Bound chromatin was eluted using stop buffer (20 mM Tris, pH 8.0; 100 mM NaCl; 20 mM EDTA; and 1% SDS) at 75 °C for 15 minutes. Eluates were incubated at 75 °C overnight to reverse cross-links, followed by treatment with proteinase K (2 μL of 10 mg/mL) at 50 °C for 6 hours and RNase A (2 μL of 10 mg/mL) at 42 °C for 2 hours. DNA was purified using the MinElute PCR Purification Kit (Qiagen, 28004) according to the manufacturer’s instructions and eluted in 12 μL of elution buffer. Purified ChIP DNA was quantified using Qubit fluorometric quantification prior to library preparation. Sequencing libraries were prepared using the KAPA HyperPrep Kit (Roche, KK8504). Paired-end sequencing was performed on an Illumina platform at Taiwan Genomic Industry Alliance Inc.

### ChIP-seq data analysis

For each ChIP-seq dataset, FASTQ files were aligned to the *Saccharomyces cerevisiae* sacCer3 reference genome (release R64-2-1) using Bowtie2 (version 2.3.5.1). Genome-wide coverage tracks were generated from aligned BAM files using bamCoverage (deepTools version 3.5.1) with a bin size of 25 bp and normalized to reads per genomic content (RPGC) to account for sequencing depth and effective genome size. Target histone modification and total H3 ChIP-seq samples were processed independently using identical parameters. To control for potential differences in nucleosome occupancy across strains and conditions, target histone modification signals were further normalized to the corresponding total H3 signals using BigWig-based normalization. The resulting H3-normalized BigWig files were used for all downstream quantitative analyses. Gene-level ChIP-seq signal quantification was performed using multiBigwigSummary (deepTools) in BED-file mode with a predefined gene list. For each gene, the average normalized ChIP-seq signal across the entire gene region was calculated, yielding one mean signal value per gene for each sample. These gene-level values were used for comparative analyses. For visualization, heatmaps were generated using deepTools based on H3-normalized ChIP-seq signals calculated across gene bodies (start codon to stop codon) with ±500 bp flanking regions. Gene-level ChIP-seq signal values from two biological replicates were averaged to generate a single representative distribution per condition. Violin plots were used to display the distribution of gene-level ChIP-seq signals. Statistical analyses, including unpaired Welch’s *t* tests, and data visualization were performed using GraphPad Prism. Principal component analysis (PCA) was performed on genome-wide ChIP-seq signal matrices generated using multiBigwigSummary in bins mode (deepTools) with a bin size of 500 bp. PCA was carried out using plotPCA (deepTools) based on RPGC-normalized BigWig files. Gene lists used in this study were generated and verified as described in fig. S3.

### *CAN1* mutation rate assay

Eleven individual colonies were randomly selected and grown overnight in YPD medium at 30 °C. Cultures were diluted 100-fold into fresh medium with or without hydroxyurea (HU) and allowed to grow to stationary phase (approximately 30 hours). Cells were subsequently diluted and plated on YPD plates to determine plating efficiency and on synthetic complete medium lacking arginine and supplemented with 60 μg/mL canavanine to select for *can1* mutants. Colony growth was scored after incubation at 30 °C for 3 days. Mutation rates were calculated using the Lea-Coulson method of the median, and statistical significance between conditions was assessed using the Mann-Whitney *U* test.

### AI-Assisted Language Editing

AI-assisted language editing was performed using ChatGPT (OpenAI, GPT-5.1). The tool was used to assist with language polishing and clarity; all scientific content, interpretation, and conclusions were generated and verified by the authors.

## Acknowledgments

We thank members of the Kao and Lo laboratories for constructive discussions. S.Y.C.’s postdoctoral fellowship (2021-2022) was supported by Academia Sinica. We thank Shin-Yi Du for technical support with flow cytometry analysis. We also thank the Biochemistry Core Facility, Institute of Cellular and Organismic Biology, Academia Sinica, for access to the homogenizer, flow cytometer, and membrane imaging system. We thank the DNA Sequencing Core Facility, Institute of Biomedical Sciences, Academia Sinica, for assistance with DNA sequencing. We thank our summer interns from National Ilan University, Chia Sim Tew and Jia-Sian Huang, for assistance with the mutation rate assay. We used OpenAI’s ChatGPT solely for language editing; all scientific content was generated, verified, and approved by the authors. No AI-generated figures were used.

## Funding

National Science and Technology Council, Taiwan (NSTC 112-2320-B-002-015-MY3 to YCL)

National Science and Technology Council, Taiwan (NSTC 113-2320-B-001-026 to CFK)

Academia Sinica (AS-IV-114-L07 to CFK)

## Author contributions

Conceptualization: SYC, YCL, CFK

Methodology: SYC, YLC, YTH, CLH, TML

Investigation: SYC, YCL, CFK

Visualization: SYC, CFK

Supervision: SYC, YCL, CFK

Writing-original draft: SYC, YCL, CFK

Writing-review & editing: SYC, YCL, CFK

## Data and materials availability

The raw sequencing data have been deposited in the NCBI Sequence Read Archive (SRA) under BioProject accession number PRJNA1399756 (individual run accessions SRR36722469-SRR36722532). Processed data have been deposited in the NCBI Gene Expression Omnibus (GEO) under accession number GSE315970.

---

**Fig. S1.**
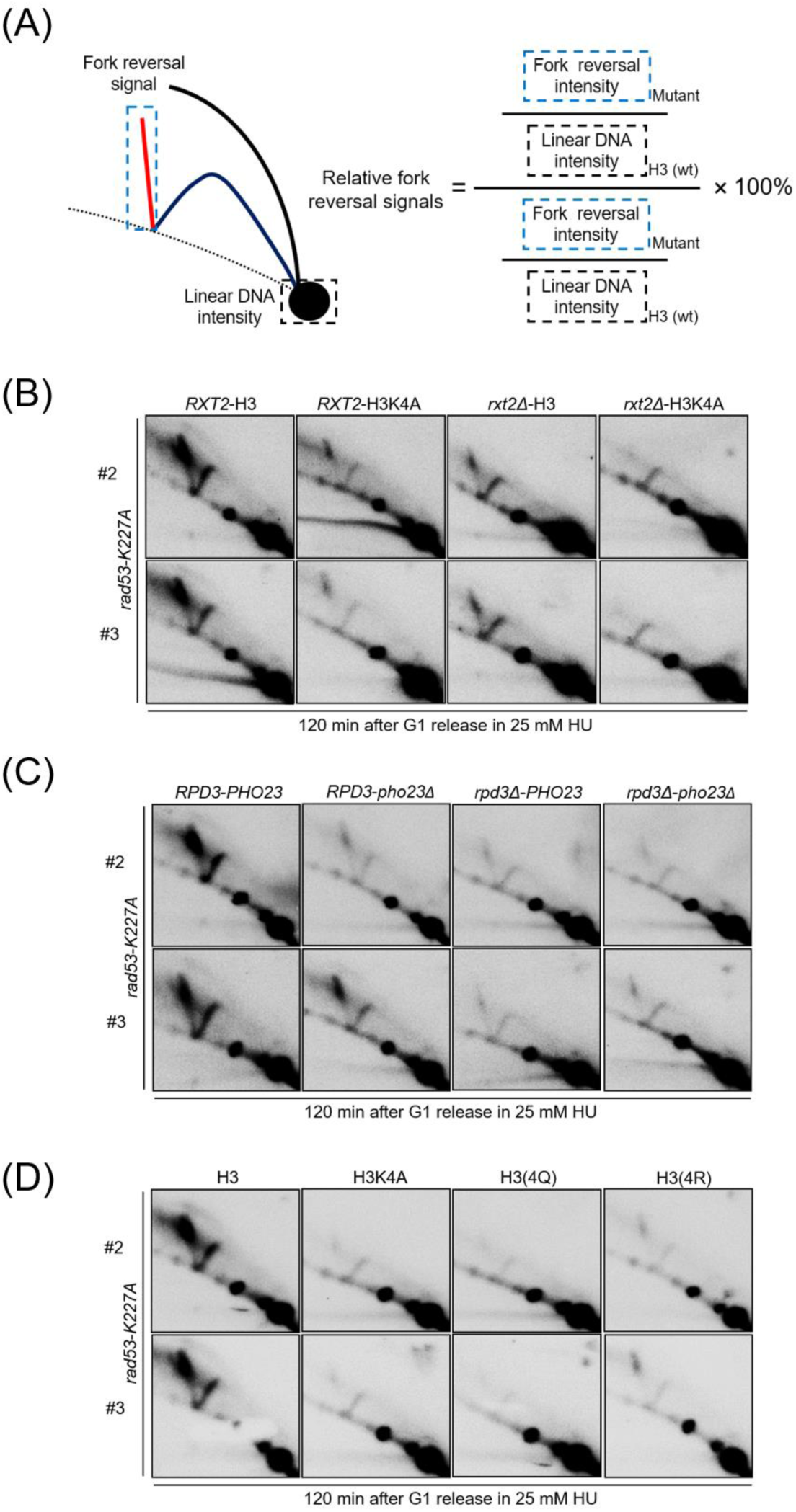
Replicate 2D gel analyses and quantification strategy. Related to Fig. 2 and Fig. 3. **(A)** Schematic illustrating the quantification of replication fork reversal signals from 2D gel images, calculated as the ratio of fork reversal signal intensity to linear DNA intensity and normalized to the corresponding H3 control. **(B-D)** Representative biological replicates of 2D gel analyses shown in Fig. 2 (B-C) and Fig. 3 (F). Replication intermediates were analyzed at the indicated loci 120 min after G1 release into 25 mM HU in *rad53-K227A* cells carrying the indicated genotypes. Each panel shows independent experimental repeats (#2 and #3), confirming the reproducibility of the observed fork reversal phenotypes.

**Fig. S2.**
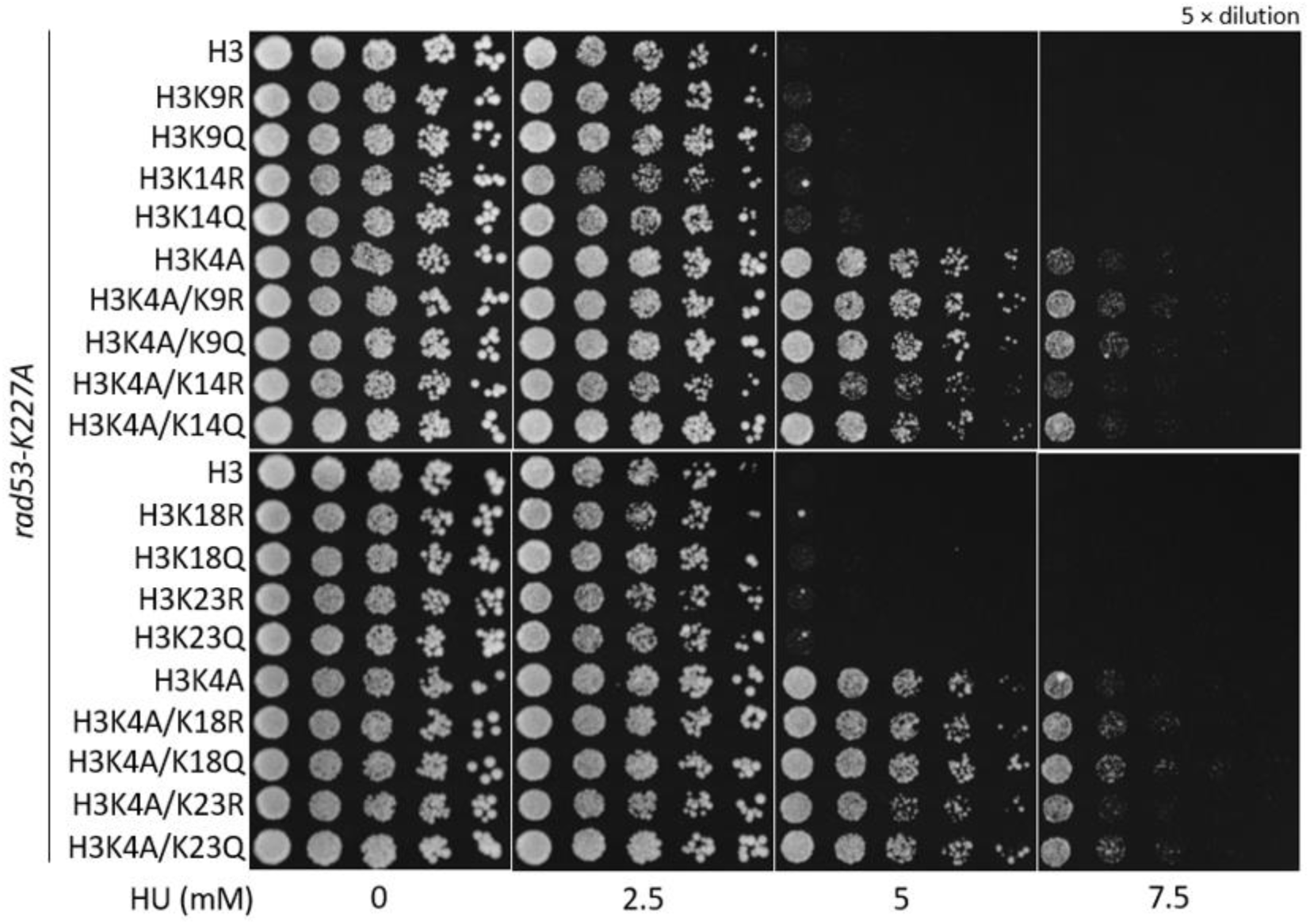
Single lysine substitutions at H3 acetylation sites do not significantly alter HU sensitivity in rad53-K227A cells. Spot dilution assays showing the growth of *rad53-K227A* cells carrying single lysine-to-glutamine (K→Q) or lysine-to-arginine (K→R) substitutions at individual histone H3 residues (K9, K14, K18, and K23), either alone or in combination with H3K4A, under increasing concentrations of hydroxyurea (HU). Serial fivefold dilutions were spotted onto YPD plates containing the indicated HU concentrations. Single-site substitutions alone did not cause obvious changes in HU sensitivity compared with the H3 control, indicating that no single acetylation site is sufficient to account for the suppression phenotype observed with combined acetylation-mimicking mutants.

**Fig. S3.**
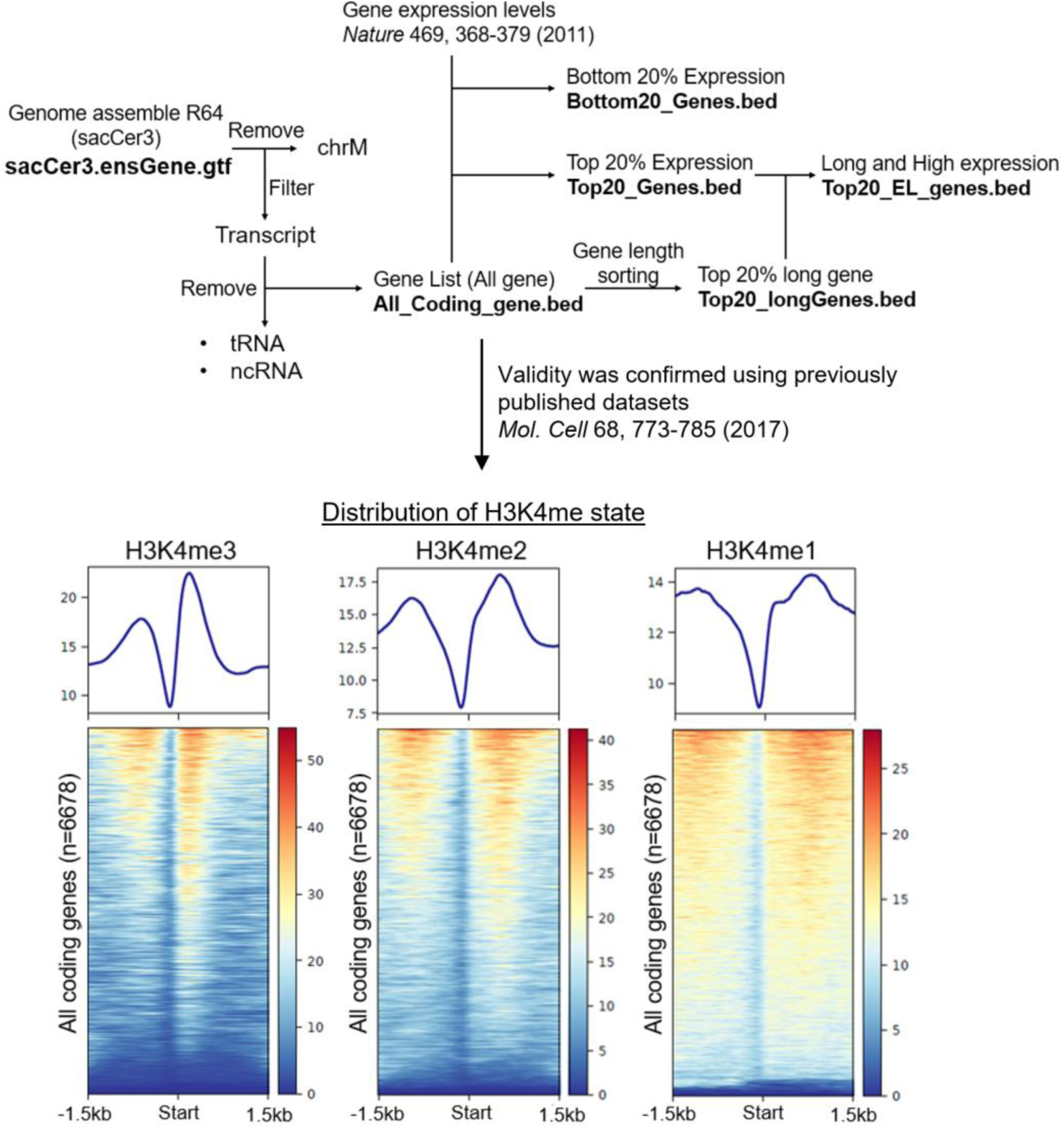
Pipeline for gene list generation and validation used in this study. Flowchart showing the procedure for generating gene lists used in ChIP-seq analyses. Gene annotations were obtained from the *Saccharomyces cerevisiae* reference genome (sacCer3, assemble R64), and protein-coding genes were selected after removing tRNA and noncoding RNA genes. Genes were classified based on expression levels (top 20% and bottom 20%) using published RNA-seq data (*Nature* 469, 368-373, 2011) and by gene length (top 20% longest genes). Gene lists were validated using previously published ChIP-seq datasets (*Mol. Cell* 68, 773-785, 2017). Representative metagene profiles and heatmaps of H3K4me1, H3K4me2, and H3K4me3 are shown.

**Fig. S4.**
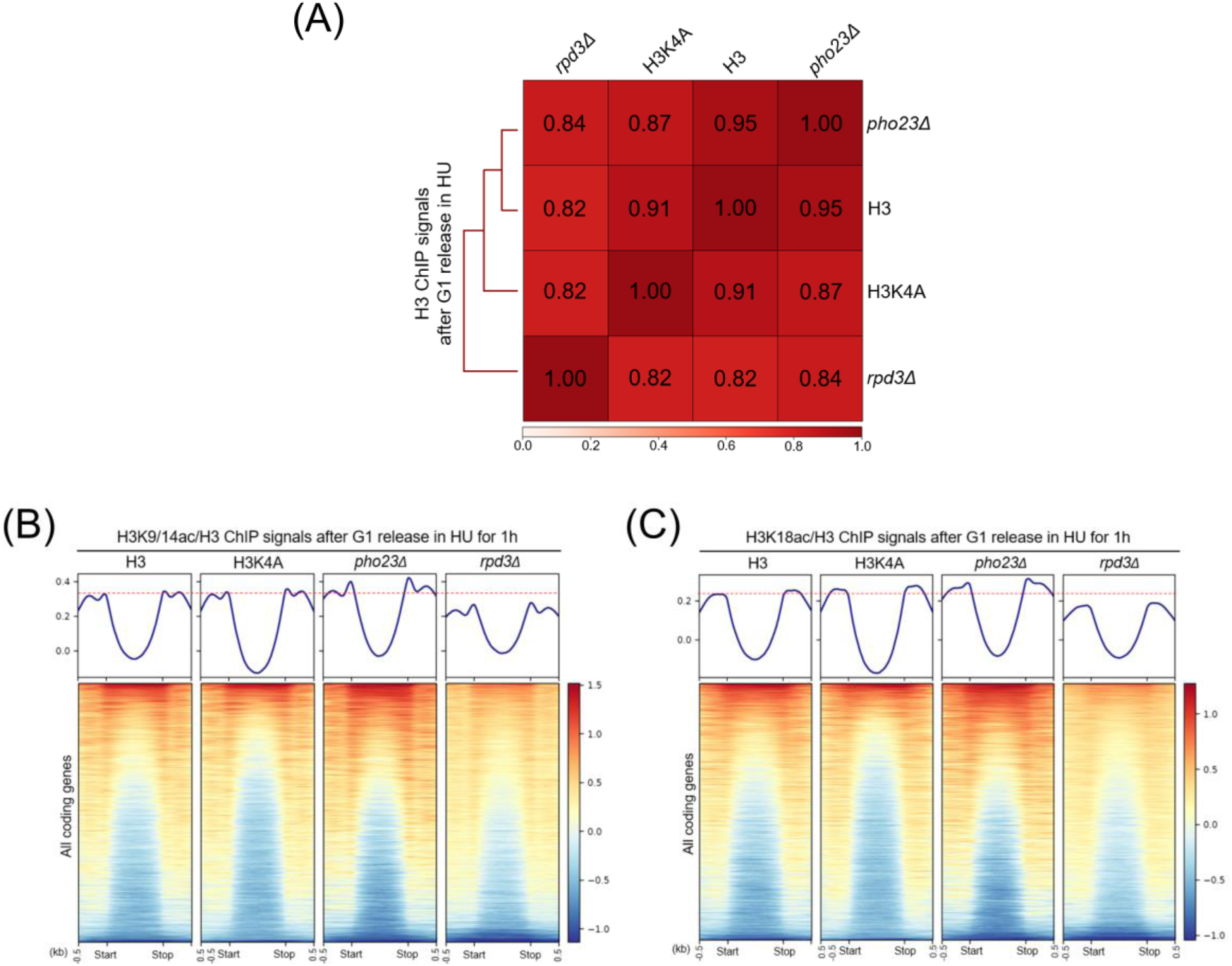
Similarity of H3 ChIP-seq profiles across strains and genome-wide distributions of H3 acetylation marks. Related to Fig. 4. **(A)** Pairwise correlation heatmap of genome-wide H3 ChIP-seq signals among the indicated strains after G1 release in HU, showing overall similarity across samples. **(B** and **C)** Metagene profiles (upper) and heatmaps (lower) of H3K9/14ac and H3K18ac ChIP-seq signals normalized to total H3 across all genes after G1 release in HU for 1 h in the indicated strains.

**Fig. S5.**
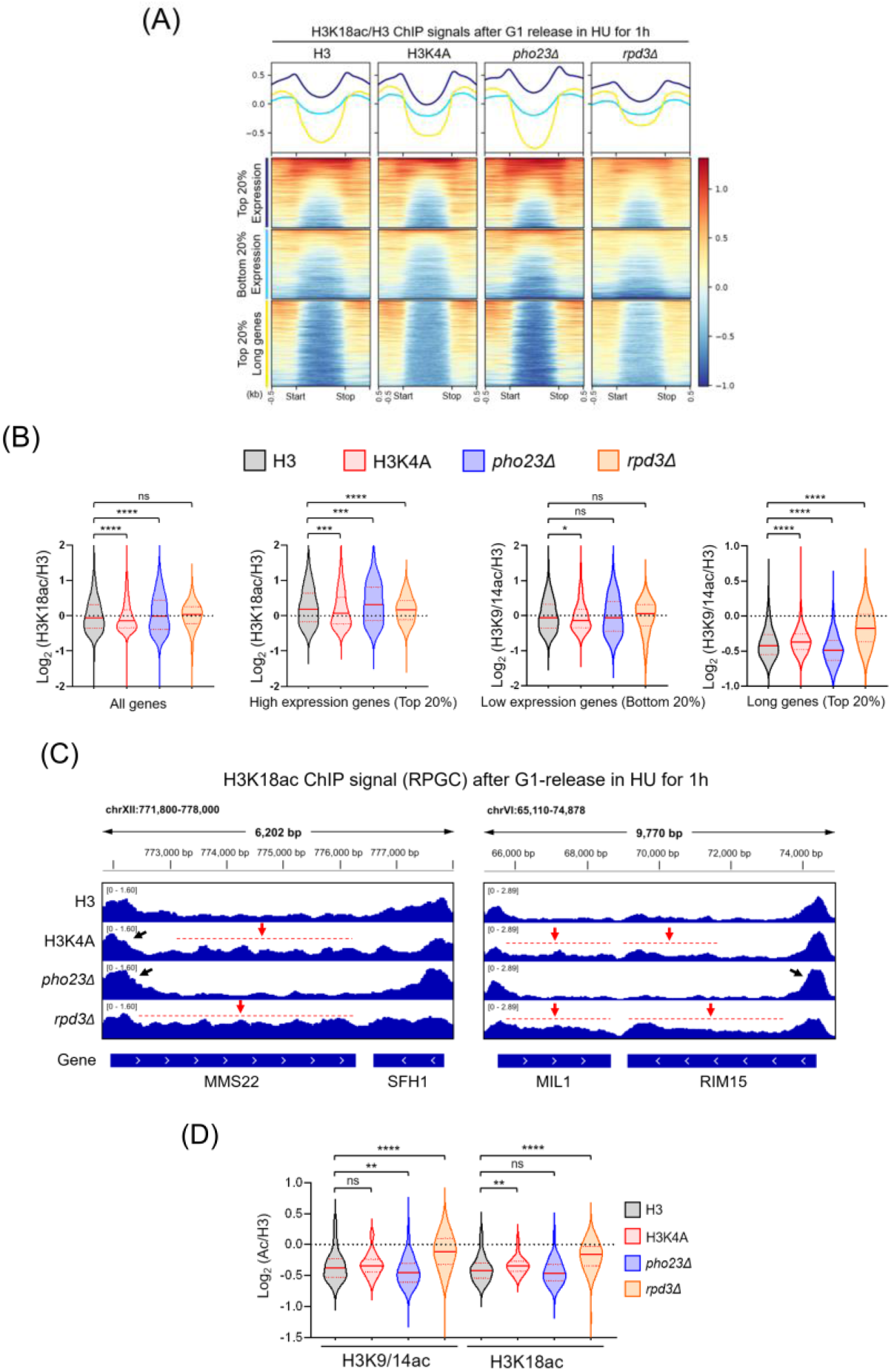
Genome-wide analysis of H3K18 acetylation following G1 release under HU treatment. Related to Fig. 4. **(A)** Metagene profiles (upper) and heatmaps (lower) showing H3K18ac ChIP-seq signals normalized to total H3 across gene bodies after G1 release in HU for 1 h in the indicated strains (H3, H3K4A, *pho23Δ*, and *rpd3Δ*). Genes are grouped by expression level (top 20% and bottom 20%) and by gene length (top 20% longest genes). **(B)** Violin plots showing genome-wide H3K18ac/H3 ChIP-seq signals across whole gene bodies for all genes, highly expressed genes (top 20%), lowly expressed genes (bottom 20%), and long genes (top 20%) in the indicated strains. **(C)** Representative genome browser views of H3K18ac ChIP-seq signals (RPGC normalized) after G1 release in HU for 1 h at the indicated loci. **(D)** Comparison of genome-wide H3K9/14ac and H3K18ac ChIP-seq signals across gene bodies of long and highly transcribed genes in the indicated strains. Statistical significance is indicated as ns (not significant), * *P* < 0.05, ** *P* < 0.01, \*\**P* < 0.001, and \*\*\**P* < 0.0001.

**Fig. S6.**
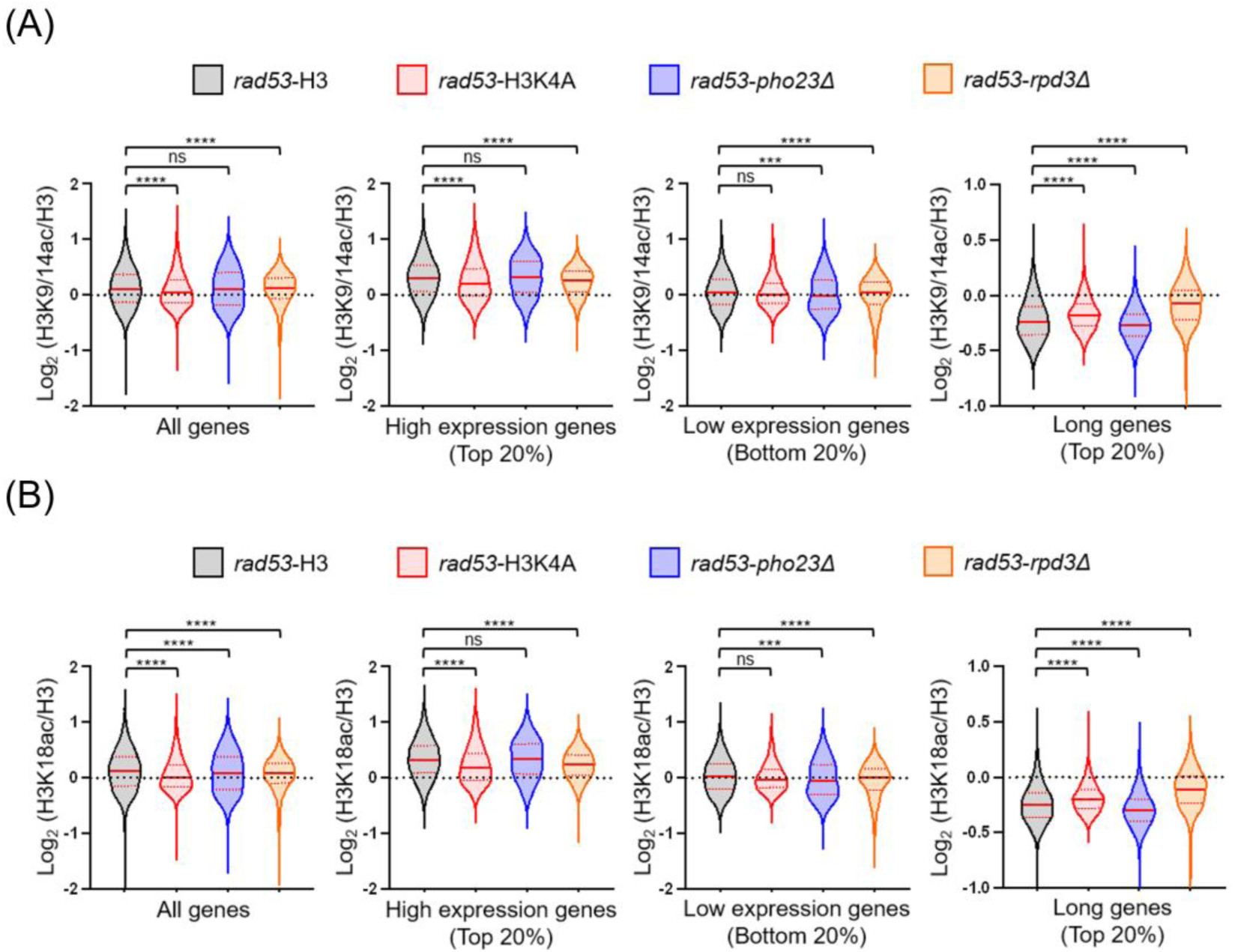
Genome-wide H3 acetylation profiles in *rad53-K227A* cells. Related to Fig. 4. Violin plots showing genome-wide **(A)** H3K9/14ac and **(B)** H3K18ac ChIP-seq signals normalized to total H3 across whole gene bodies in *rad53-K227A* cells carrying the indicated genotypes (H3, H3K4A, *pho23Δ*, and *rpd3Δ*). Data are shown for all genes, highly expressed genes (top 20%), lowly expressed genes (bottom 20%), and long genes (top 20%). Statistical significance is indicated as ns (not significant),* *P* < 0.05, ** *P* < 0.01, \*\**P* < 0.001, and \*\*\**P* < 0.0001.

**Supplementary Table 1:**
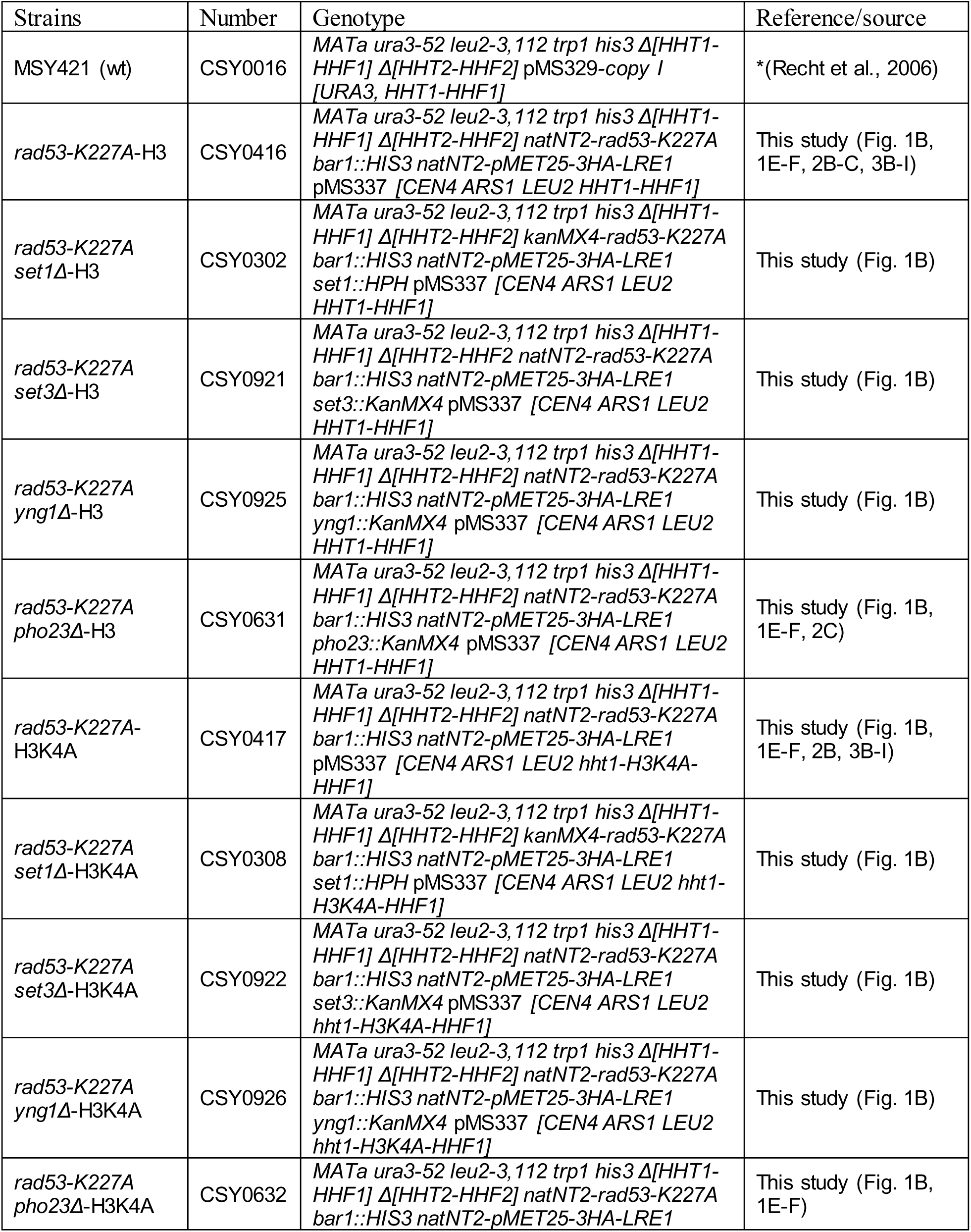
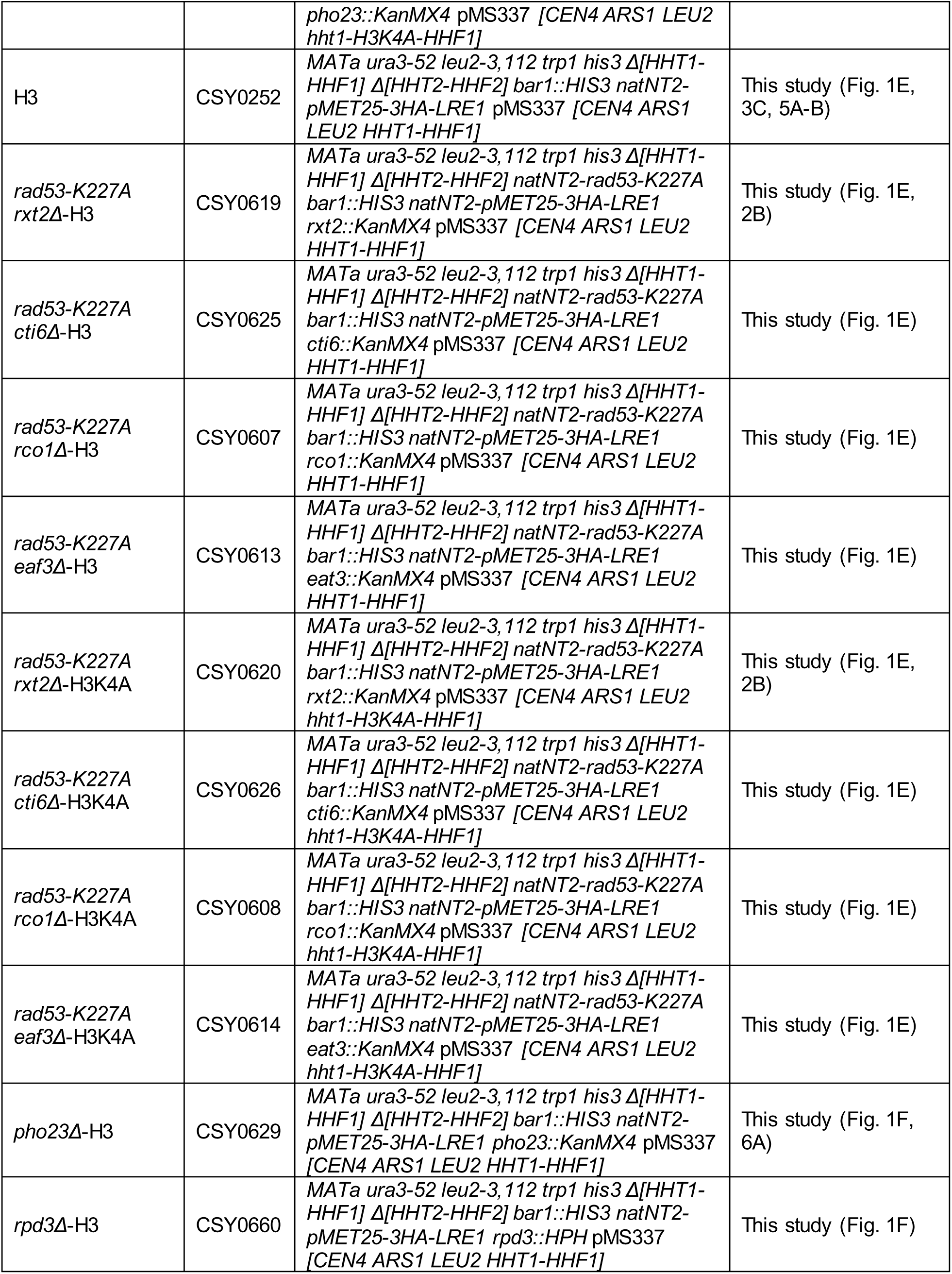
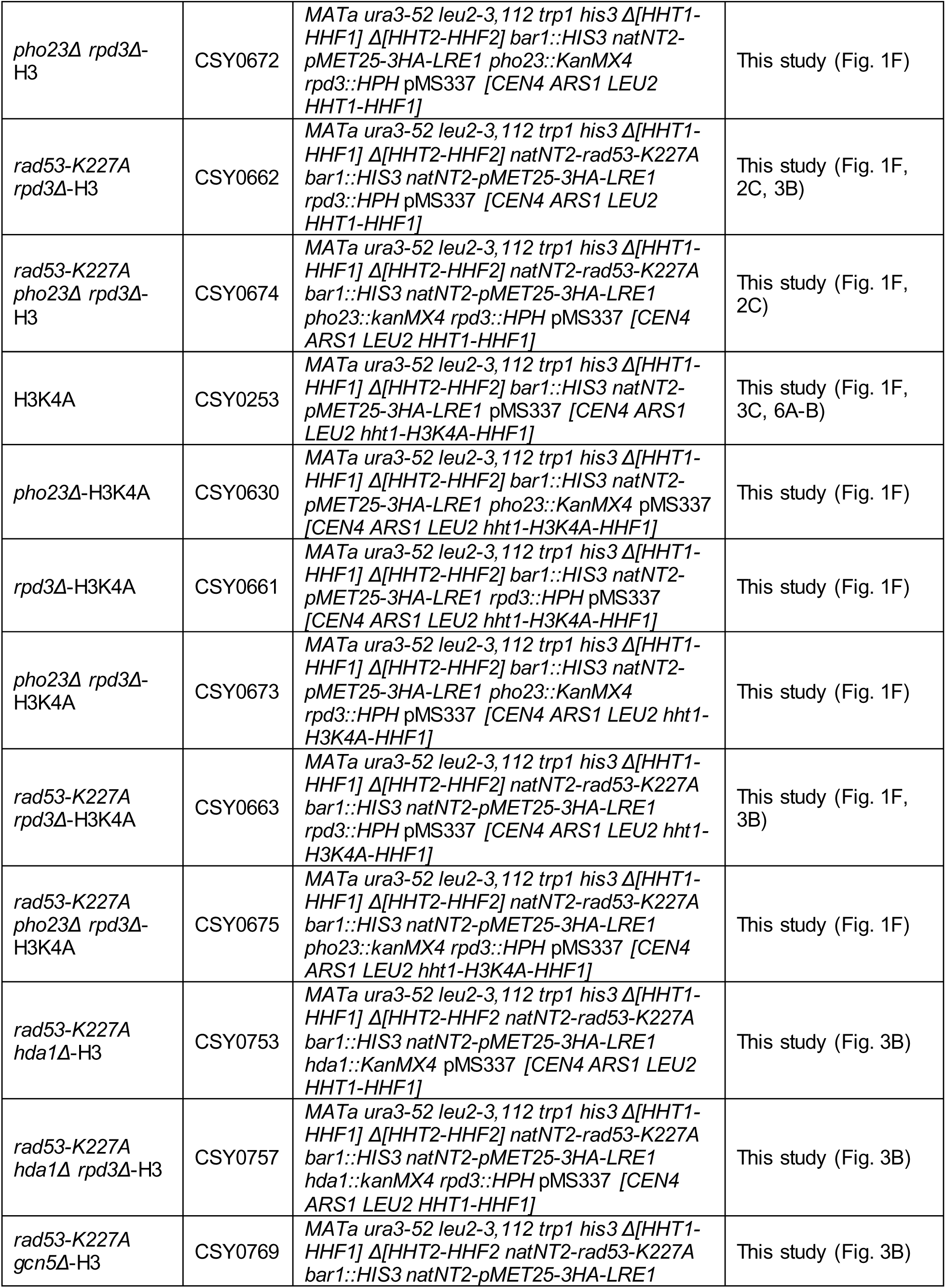
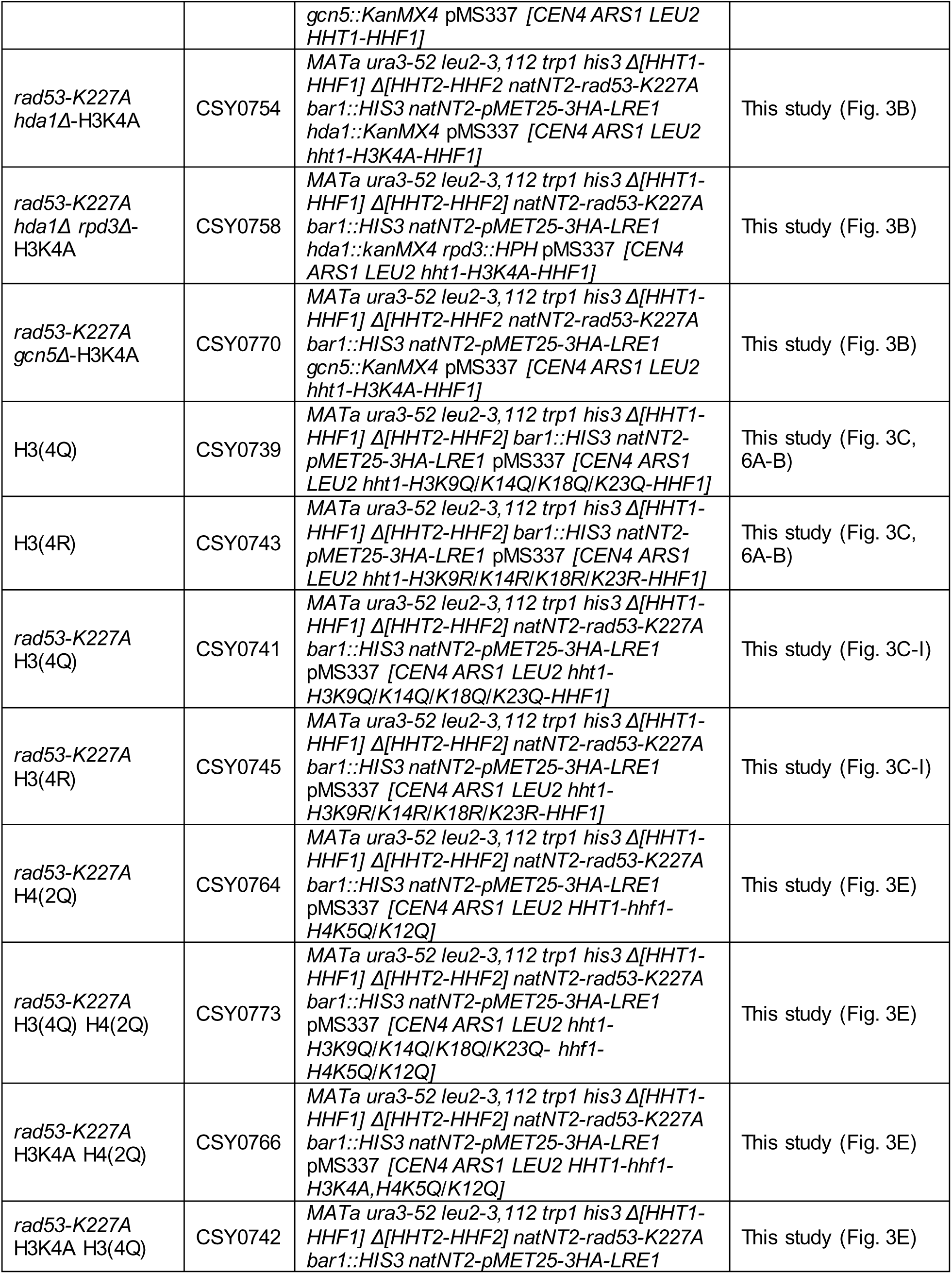
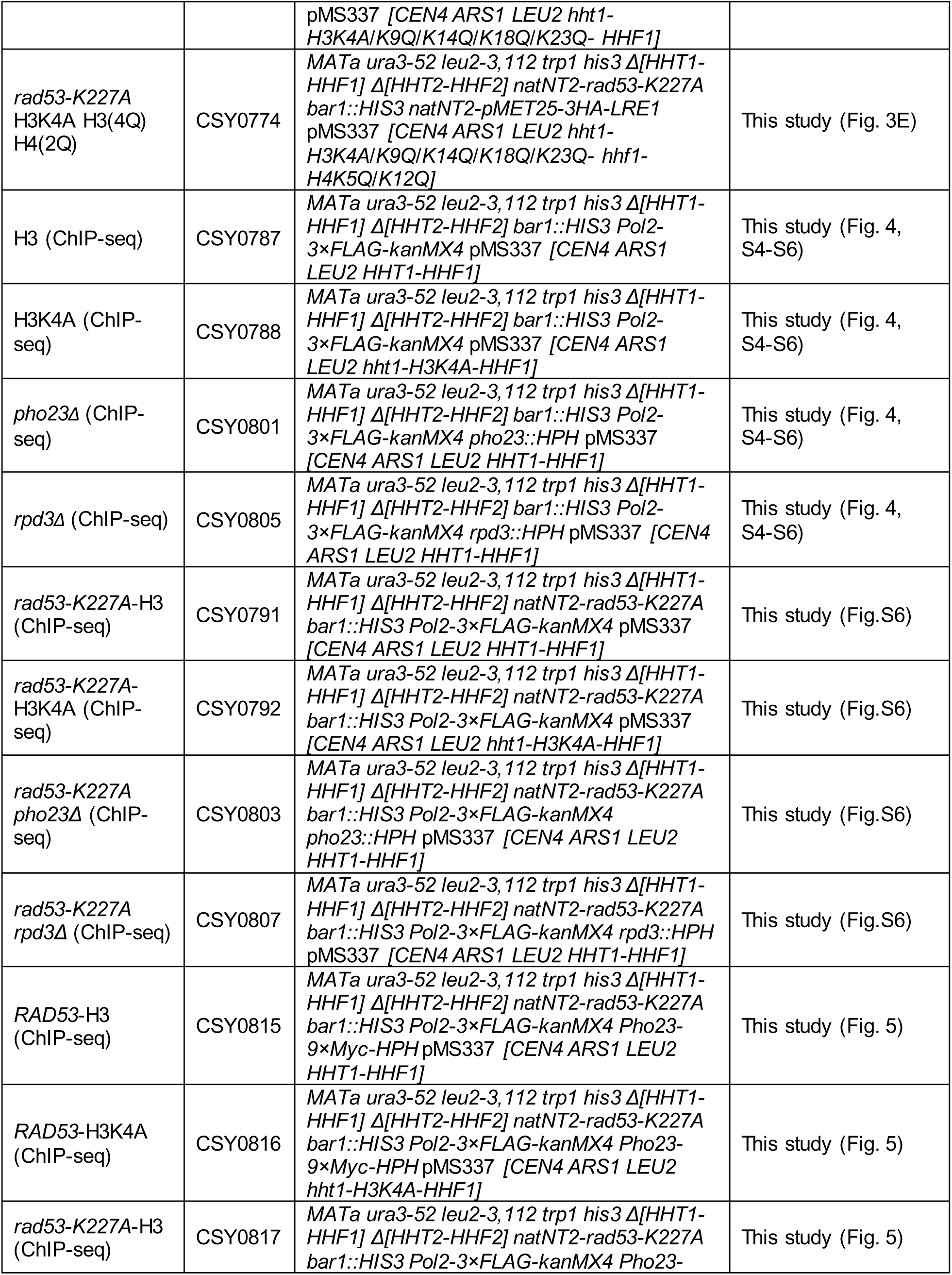
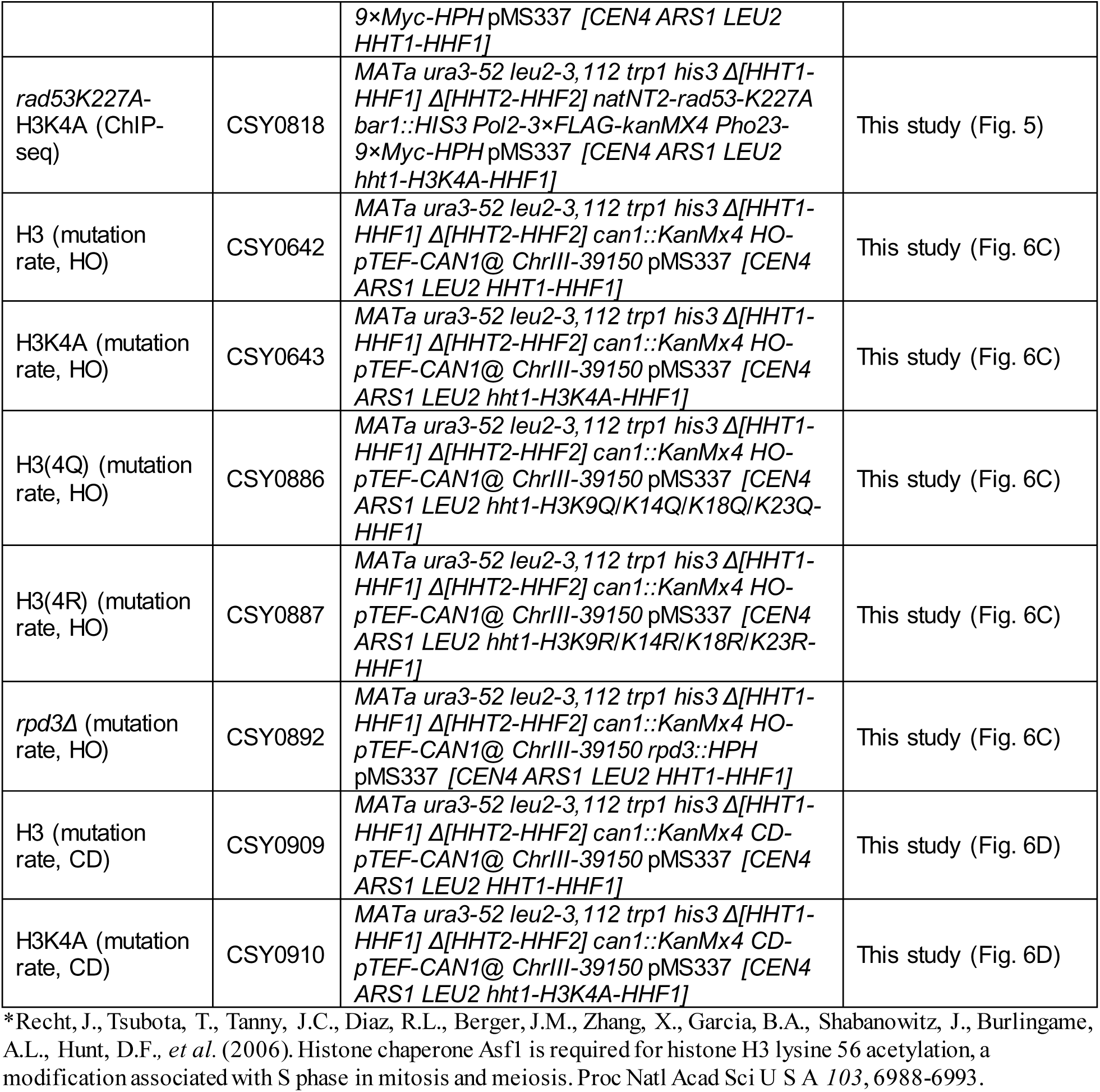
Strains used in this study.

**Supplementary Table 2:**
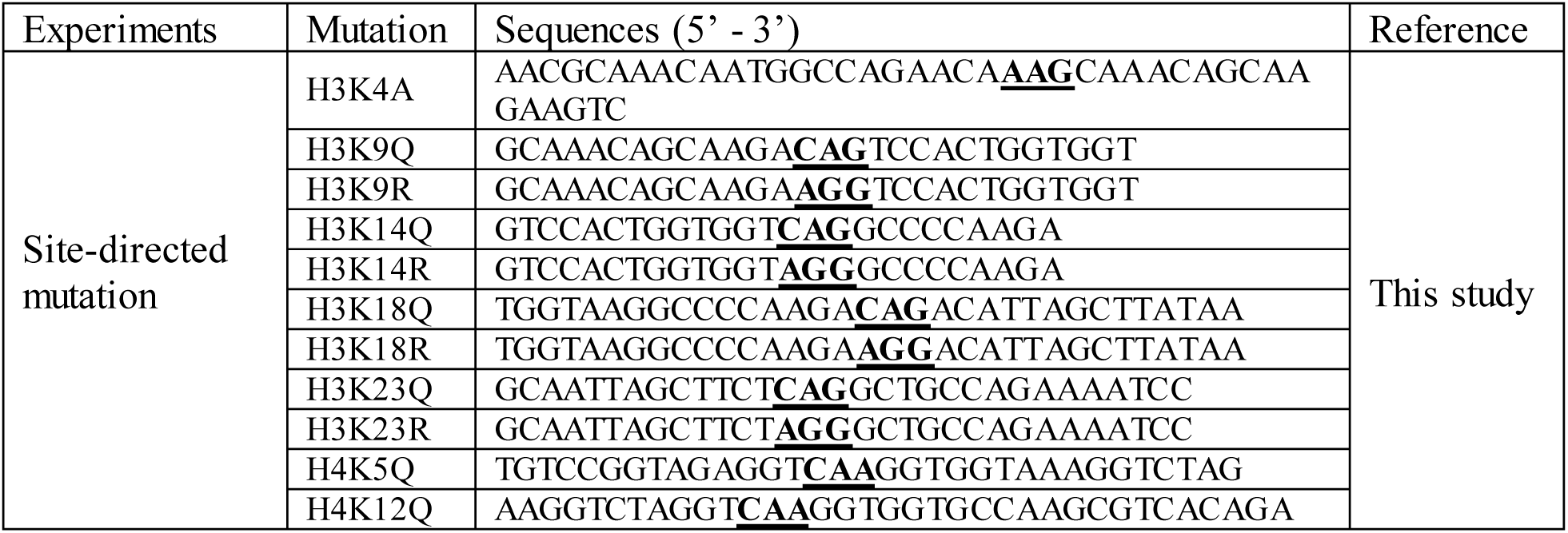
Primers used for site-direct mutagenesis in this study.

**Supplementary Table 3:**
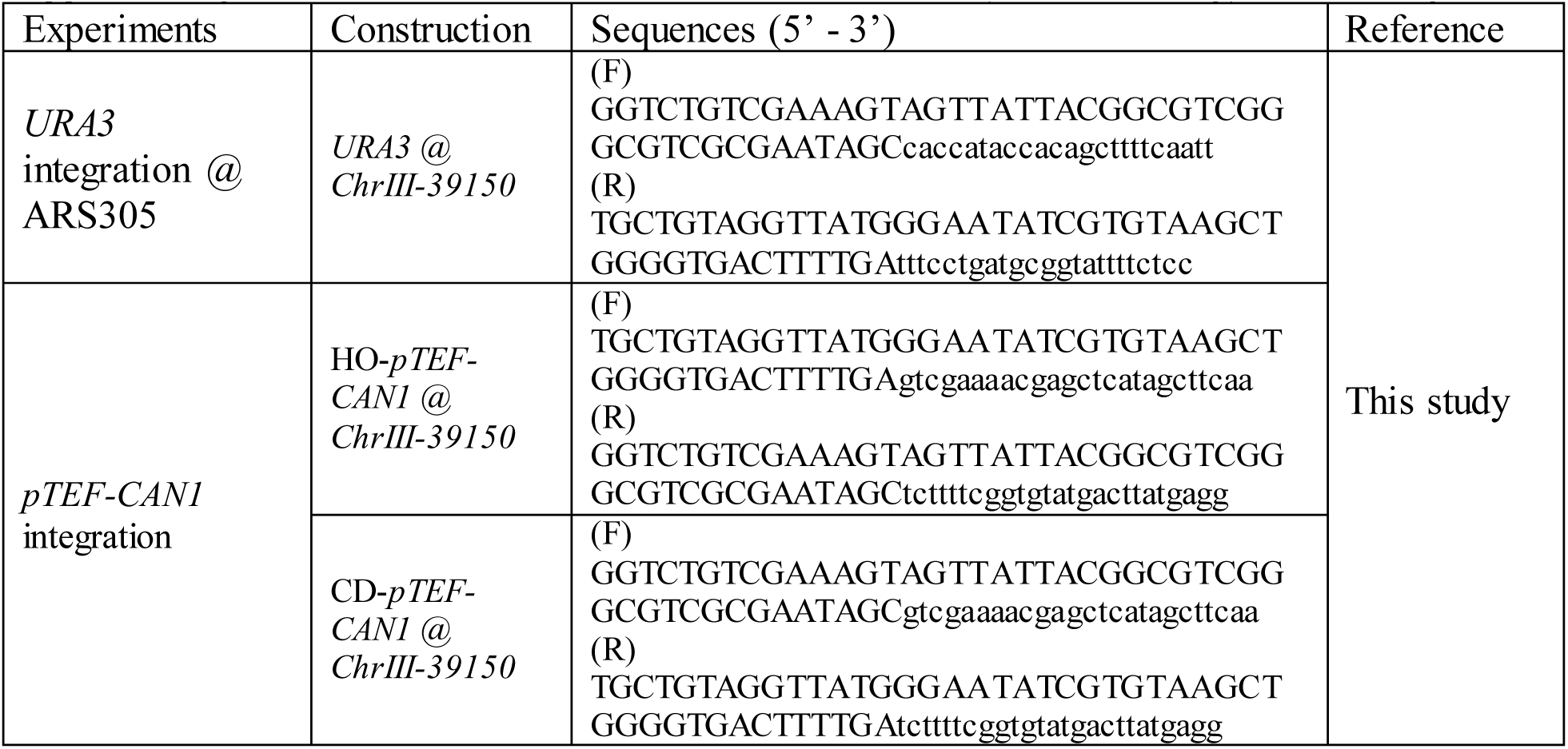
Primers used for strain construction (mutation assay) in this study.

**Supplementary Table 4:**
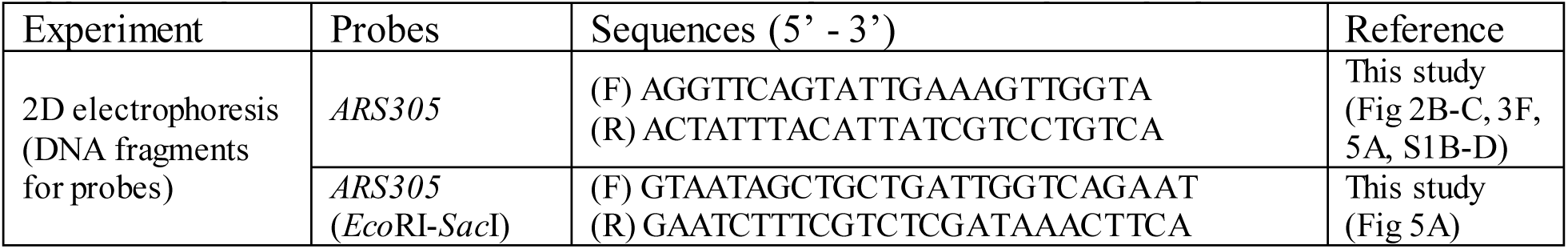
Primers used for PCR-amplification for probe preparation.

